# Mitochondrial stress signaling shapes the nuclear response to loss of the chromatin reader MRG-1

**DOI:** 10.64898/2026.08.14.744963

**Authors:** Carole Zaratiegui, Fernanda Rezende Pabst, Maria-Jesus Rodriguez Palero, Peter Meister, Marta Artal-Sanz, Daphne S. Cabianca

**Affiliations:** Institute of Functional Epigenetics, Helmholtz Munich, Oberschleißheim, Germany; LMU Faculty of Medicine, Munich, Germany; Andalusian Centre for Developmental Biology, Consejo Superior de Investigaciones Científicas/Junta de Andalucía/Universidad Pablo de Olavide, Seville, Spain; Institute of Cell Biology, University of Bern, Bern, Switzerland

## Abstract

Perinuclear sequestration of heterochromatin is a major conserved feature of nuclear architecture. In *Caenorhabditis elegans*, the euchromatic reader MRG-1 was previously shown to promote peripheral localization of heterochromatin through an indirect mechanism that remained largely unknown. Here, we show that loss of MRG-1 activates a mitochondrial stress response. Genetic ablation of the PMK-3/MAPK mitochondrial stress regulator CBP-3 reveals that this pathway contributes to both detachment of a heterochromatic reporter from the nuclear periphery and approximately one-third of the transcriptional changes induced by *mrg-1* depletion. Strikingly, loss of *cbp-3* in MRG-1-deficient animals exacerbates mitochondrial dysfunction, fertility defects and embryonic lethality, indicating that part of the nuclear response induced by MRG-1 loss contributes to adaptation to mitochondrial stress rather than constituting a defect in heterochromatin 3D organization as previously thought. Together, our findings identify mitochondrial stress signaling as an unexpected mediator of the nuclear response to MRG-1 loss, demonstrating its contribution to gene regulation while supporting the idea that stress-induced changes in cellular physiology can also shape nuclear organization.

## Introduction

In the great majority of eukaryotic cells, heterochromatin accumulates at the nuclear periphery, representing a major evolutionarily conserved feature of nuclear architecture (Solovei et al. 2016). Perinuclear sequestration of heterochromatin shapes genome function (Attar et al. 2025; Lewis et al. 2026; Marin et al. 2025), and association of individual genes with the nuclear periphery can influence their transcriptional activity (Finlan et al. 2008; Reddy et al. 2008), underscoring the importance of understanding how this spatial organization is established and regulated.

Pioneering work in *C. elegans* demonstrated that heterochromatin anchoring to the nuclear periphery requires methylation of histone H3 lysine 9 (H3K9me) (Towbin et al. 2012). This process is mediated by the chromodomain protein CEC-4, a highly specific H3K9me reader localized at the nuclear envelope (Gonzalez-Sandoval et al. 2015). However, although the H3K9me–CEC-4 pathway predominates in early embryos, in differentiated larval cells, additional H3K9me-independent mechanisms that maintain heterochromatin at the nuclear periphery are induced (Cabianca et al. 2019; Gonzalez-Sandoval et al. 2015). Likewise, the role of H3K9me2/me3 in heterochromatin anchoring is conserved in mammals (Bian et al. 2020; Harr et al. 2020; Kind et al. 2013; Marin et al. 2025), where distinct anchoring mechanisms have been reported, operating at different stages of development (Solovei et al. 2013) and across tissues (Robson et al. 2016). Nonetheless, H3K9me-independent mechanisms that regulate heterochromatin positioning remain largely unexplored.

In *C. elegans*, peripheral localization of heterochromatin in intestinal cells of larvae requires the chromatin reader MRG-1, which acts in parallel to the CEC-4/H3K9me pathway (Cabianca et al. 2019). MRG-1 and its homologues – yeast Eaf3 and human MRG15 – are enriched at H3K36me3-marked regions (Cabianca et al. 2019; Guan et al. 2023; Zhang et al. 2006). As in other species, H3K36me3 is associated with actively transcribed gene bodies in *C. elegans* (Liu et al. 2011). Accordingly, MRG-1 is largely excluded from heterochromatic domains (Cabianca et al. 2019), indicating that it regulates heterochromatin positioning through an indirect mechanism. However, the pathways involved remain unknown.

Here, using MRG-1-dependent heterochromatin positioning as a model, we identify mitochondrial stress signaling as a mediator of MRG-1-dependent nuclear changes. We further show that blocking the stress response partially restores nuclear phenotypes while compromising organismal fitness, suggesting that MRG-1-dependent gene expression and 3D chromatin changes are, at least in part, aspects of an adaptive response to mitochondrial stress.

## Results

### Mitochondrial stress signaling contributes to MRG-1 loss-induced detachment of a heterochromatic reporter from the nuclear periphery

To gain insight into the pathways affected by MRG-1 loss, we analyzed RNA-seq data from *mrg-1* knockout L1 larvae (Cabianca et al. 2019). Genes upregulated in *mrg-1* mutants were enriched for GO categories related to innate immunity and defense against bacterial infection (Fig. 1A), transcriptional programs that are characteristic of mitochondrial stress responses (Kwon et al. 2018; Liu et al. 2014; Pellegrino et al. 2014). In addition, we identified six established markers of mitochondrial stress among the upregulated genes (Campos et al. 2021; Munkácsy et al. 2016) (Supplemental Fig. S1A), whereas six downregulated genes encode mitochondrial proteins (Supplemental Fig. S1B). Together, these observations indicate that loss of MRG-1 induces a transcriptional signature consistent with altered mitochondrial homeostasis and activation of mitochondrial stress signaling. We therefore asked whether mitochondrial stress signaling contributes to the nuclear phenotypes caused by loss of MRG-1.

**Fig. 1.**
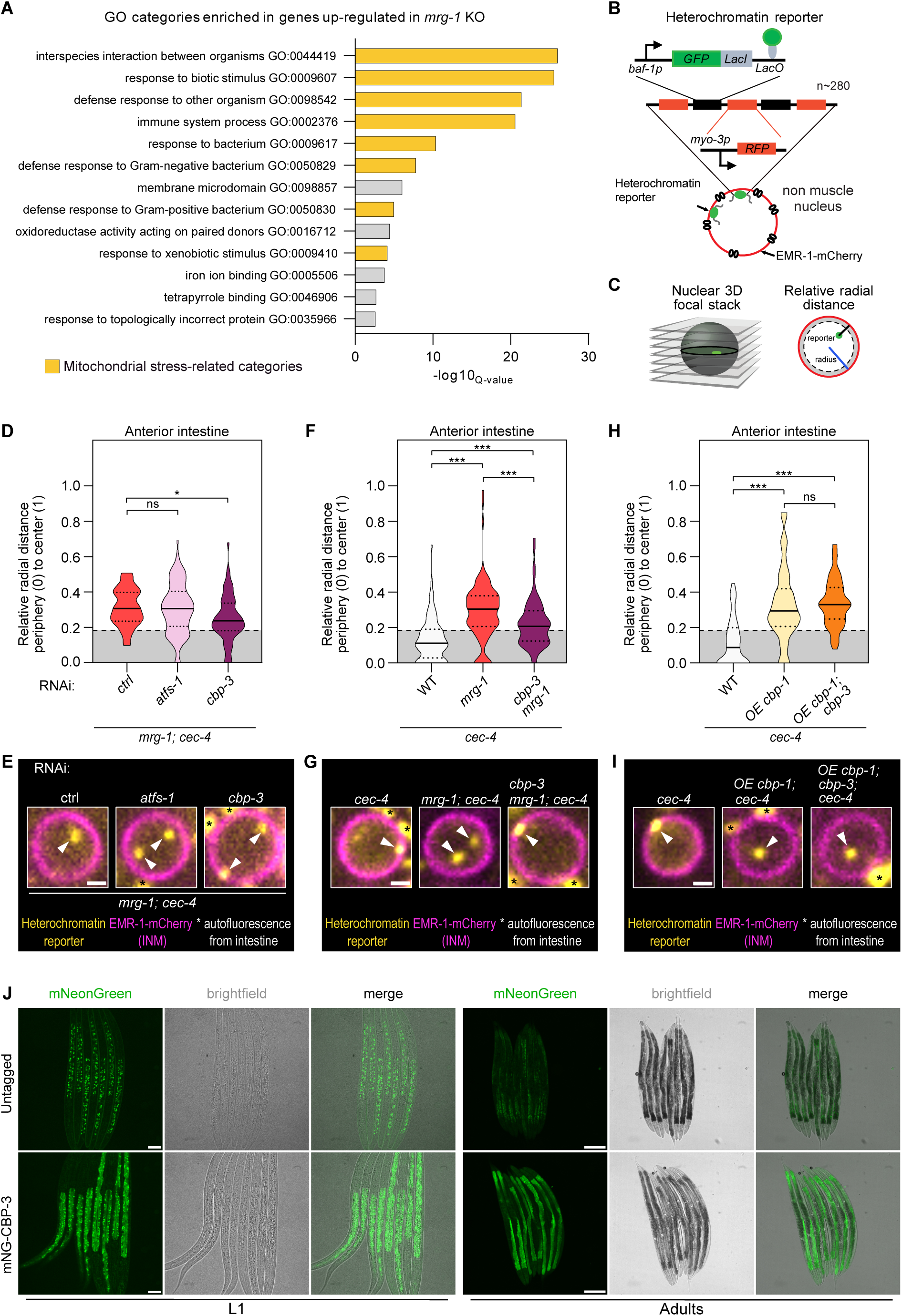
Mitochondrial stress signaling contributes to detachment of a heterochromatic reporter from the nuclear periphery following loss of MRG-1. **A)** Gene Ontology (GO) enrichment analysis of genes upregulated in *mrg-1* knockout L1 larvae (Cabianca et al., 2019). Mitochondrial stress-related GO categories are highlighted in yellow. **B)** Schematic representation of the heterochromatin reporter. **C)** Strategy for quantifying the heterochromatin reporter positioning. The shortest distance between each reporter focus and the nuclear periphery was normalized to the radius of each intestinal nucleus (relative radial distance). Reporter distribution is shown relative to the most peripheral third of the nucleus area (grey zone). **D)** Violin plot showing the relative radial distance distribution of the reporter foci in anterior intestinal nuclei of *mrg-1; cec-4* animals exposed to the indicated RNAi. Relative radial distances range from the nuclear periphery (0) to the nuclear center (1). The most peripheral third of the area of the nucleus is highlighted in grey. The thick horizontal line indicates the median, while the dotted lines represent the first (lower one) and third (higher one) quartiles. Results are from 3 biological replicates. **E)** Single focal planes of representative L1 larvae expressing the heterochromatin reporter and EMR-1–mCherry in the genotypes and RNAi conditions shown in (D). Insets show an enlarged anterior intestinal nucleus. INM: inner nuclear membrane. **F)** Same as (D), but for the indicated mutant genotypes. Results are from 4 biological replicates. **G)** Representative images as in (E), but corresponding to the mutant genotypes shown in (F). H) Same as (D), but for the indicated genotypes. OE denotes intestine-specific overexpression of CBP-1 (*ges-1p::cbp-1*). Results are from 3 biological replicates. I) Representative images as in (E), but corresponding to the mutant genotypes shown in (H). J) Single focal planes of representative L1 larvae (left) or adults (right) expressing neonGreen-CBP-3 from *cbp-3* endogenous locus or untagged WT animals as control for background autofluorescence. For E, G and I, scale bar = 1 µm. For J, scale bar L1s = 20 µm; adults = 200 µm. For D, statistical significance was assessed using one-way parametric ANOVA. For F and H, statistical significance was assessed using a Kruskal–Wallis non-parametric ANOVA. Pairwise comparisons were performed as indicated in each panel. *p< 0.05, ***p < 0.001, ns = not significant. For D, F, H, n (foci scored per condition) ≥ 154. Exact p-values and n (foci scored per condition) values are provided in Supplemental Table ST3.

To address this question, we focused on the transcription factor ATFS-1, which drives the canonical mitochondrial unfolded protein response (UPRmt) in *C. elegans* (Haynes and Ron 2010; Nargund et al. 2012, 2015) and on CBP-3, a key component of the PMK-3/MAPK mitochondrial stress pathway required for a transcriptional response to mitochondrial stress that is independent of ATFS-1 (Munkácsy et al. 2016). Specifically, we used RNAi to individually knock down *atfs-1* and *cbp-3* and monitor the localization of a well-established *lacO*/GFP-LacI-based reporter for heterochromatin positioning (Cabianca et al. 2019; Gonzalez-Sandoval et al. 2015; Meister et al. 2010b; Towbin et al. 2012), which was instrumental in identifying both CEC-4 and MRG-1 (Cabianca et al. 2019; Gonzalez-Sandoval et al. 2015) (Fig. 1B).

Reporter position was quantified by measuring the shortest distance between the reporter and the inner nuclear membrane (INM), abelled by EMR-1/Emerin-mCherry, relative to the nuclear radius (Meister et al. 2010a) (Fig. 1C), in the four anterior intestinal cells (Supplemental Fig. S1C), where delocalization is most pronounced (Cabianca et al. 2019). Because both *mrg-1* and *cec-4* are required for complete reporter detachment (Cabianca et al. 2019), the experiments were performed in this sensitized background.

As previously reported (Cabianca et al. 2019), the heterochromatin reporter is completely detached from the nuclear periphery in anterior intestinal cells of *mrg-1; cec-4* double mutants, with the median distribution falling outside the outermost nuclear region, and this phenotype was unaffected by control RNAi (Fig. 1D and 1E). We found that knockdown of *atfs-1* did not affect reporter positioning (Fig. 1D and 1E). In contrast, depletion of *cbp-3* partially restored perinuclear localization (Fig. 1D and 1E), an effect confirmed using a *cbp-3* mutant allele (Fig. 1F and 1G). These results identify CBP-3 as a mediator of heterochromatin reporter detachment from the nuclear periphery following loss of MRG-1.

It was previously shown that the CBP/p300 histone acetyltransferase CBP-1 is required for the detachment of the heterochromatic reporter in *mrg-1; cec-4* double mutants and that its overexpression is sufficient to induce reporter detachment in *cec-4* mutants, thereby phenocopying the loss of *mrg-1* (Cabianca et al. 2019). Because both CBP-1 and CBP-3 are required for reporter detachment following MRG-1 loss, we next asked whether CBP-3 functions upstream or downstream of CBP-1. To address this, we tested whether loss of CBP-3 could suppress the reporter detachment induced by CBP-1 overexpression. In contrast to its effect in *mrg-1; cec-4* mutants, CBP-3 depletion did not suppress detachment of the heterochromatic reporter caused by CBP-1 overexpression (Fig. 1H and 1I), consistent with CBP-3 acting downstream of MRG-1 and upstream of, or in parallel to, CBP-1 to promote reporter detachment from the nuclear periphery. The domain architecture of CBP-3 further supports this interpretation. As a truncated paralog of CBP-1, it lacks the catalytic histone acetyltransferase and bromodomain domains, while retaining only a zinc finger domain (Supplemental Fig. S1D), making it unlikely that CBP-3 directly modifies chromatin.

To determine the expression pattern of CBP-3, we generated an endogenously tagged fluorescent allele using CRISPR/Cas9 and found that CBP-3 is expressed strongly in the intestine of both L1 larvae and adults (Fig. 1J), consistent with its role in regulating positioning of the heterochromatic reporter in this tissue.

### Mitochondrial stress is sufficient to induce heterochromatin reporter detachment in a CBP-3-dependent manner

The requirement for CBP-3 for reporter detachment following loss of MRG-1 indicated that a mitochondrial stress pathway is involved in this phenotype. We therefore asked whether mitochondrial dysfunction itself is sufficient to induce delocalization. To test this, we directly perturbed mitochondrial function and monitored the localization of the heterochromatin reporter. In intestinal cells of L1 larvae, complete detachment of the reporter from the nuclear periphery requires loss of both CEC-4 and MRG-1, as these pathways redundantly anchor the heterochromatic reporter (Cabianca et al. 2019). We therefore used *cec-4* single mutants as a sensitized background and induced mitochondria dysfunction by RNAi-mediated depletion of components of the electron transport chain (ETC; *nuo-1* and *nuo-6* for complex I, *sdha-1* and *sdhd-1* for complex II, *cyc-1* and *F45H10.2* for complex III, *cco-1* and *cco-4* for complex IV) and ATP synthase (complex V; *atp-3* and *asg-2*). Strikingly, depletion of both tested components of ETC complexes III and IV, as well as one of the two ATP synthase subunits tested (*atp-3*), caused an internalization of the heterochromatic reporter away from the nuclear periphery in *cec-4* mutants (Fig. 2A), with the median reporter position shifting beyond the outermost nuclear zone (Fig. 2A), thereby phenocopying the effect observed in *mrg-1; cec-4* double mutants. In contrast, depletion of components of ETC complexes I or II had no detectable effect on reporter positioning (Fig. 2A), indicating that not all forms of ETC dysfunction equally affect reporter positioning.

**Fig. 2.**
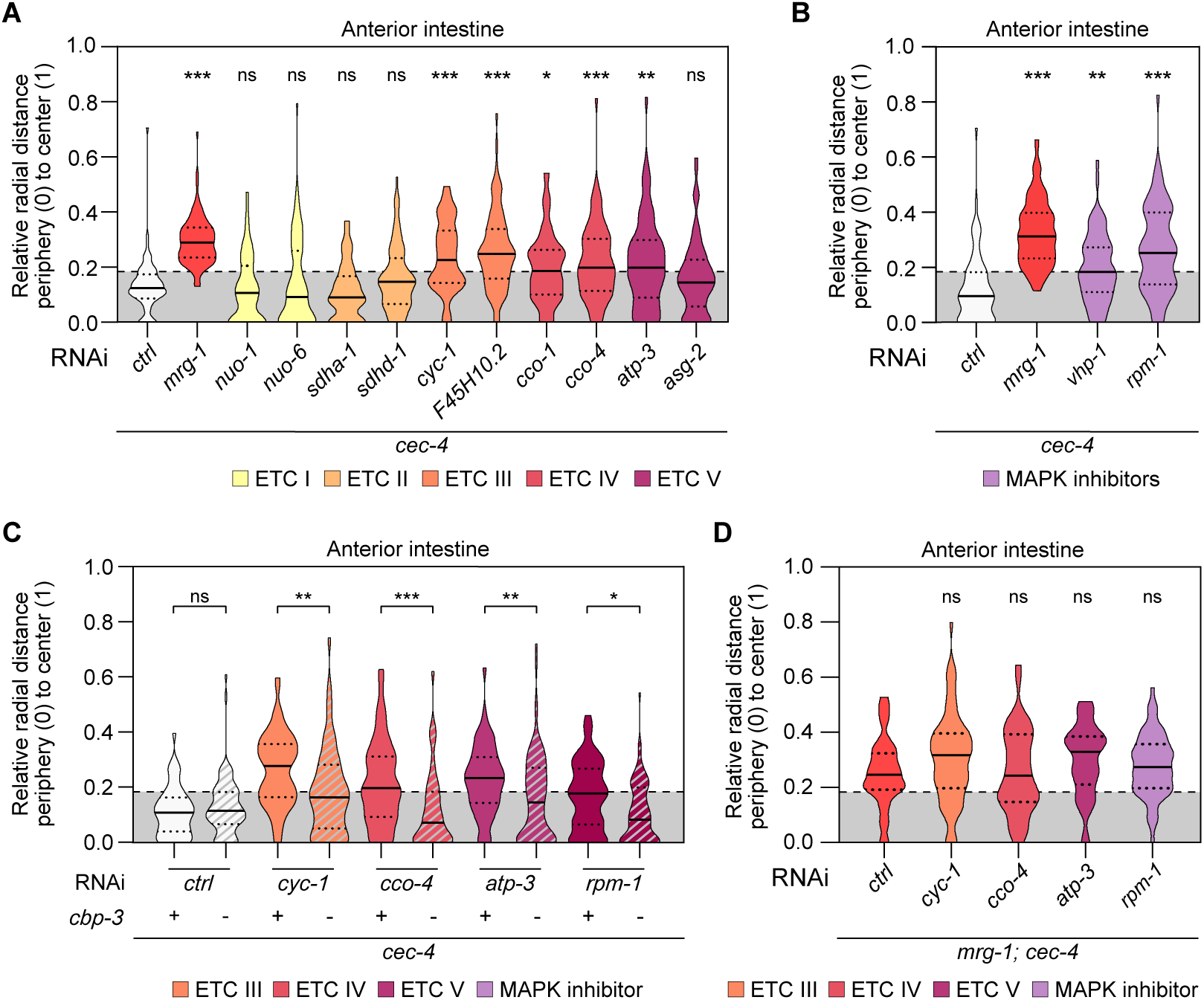
Mitochondrial dysfunction is sufficient to induce CBP-3-dependent detachment of the heterochromatic reporter from the nuclear periphery. **A)** Violin plot showing the relative radial distance distribution of the heterochromatin reporter foci in anterior intestinal nuclei of *cec-4* mutant L1s exposed to the indicated RNAi. Relative distances range from the nuclear periphery (0) to the nuclear center (1). The most peripheral third of the nucleus is highlighted in grey. The thick horizontal line indicates the median, while the dotted lines represent the first (lower one) and third (higher one) quartiles. Results are from at least 3 biological replicates. **B)** Same as (A) but for the indicated RNAi conditions. Results are from 3 biological replicates. **C)** Same as (A) but in the indicated genotypes, treated with the indicated RNAi. Results are from at least 3 biological replicates. **D)** Same as (A) but in *mrg-1; cec-4* mutants treated with the indicated RNAi. Results are from 3 biological replicates. For A, B and C, statistical significance was assessed using a Kruskal–Wallis non-parametric ANOVA. For D, statistical significance was assessed using a one-way parametric ANOVA. Pairwise comparisons were performed against the reference condition or as indicated in each panel. *p< 0.05, **p< 0.01, ***p< 0.001, ns = not significant. For A-D, n (foci scored per condition)≥ 87. Exact p-values and n (foci scored per condition) values are provided in Supplemental Table ST3.

Next, for one positive hit from each delocalizing ETC complex, we examined reporter positioning in mid-intestinal cells (Supplemental Fig. S1C). We chose this region because *mrg-1* mutants also display reporter internalization there, albeit to a lesser extent than in the anterior intestine (Cabianca et al. 2019). Knockdown of *cyc-1*, *cco-4*, or *atp-3* each induced reporter internalization in middle intestine (Supplemental Fig. S2A). However, the median reporter position remained within the outermost nuclear zone, similar to the phenotype observed upon *mrg-1* RNAi, indicating a milder defect than that seen in anterior intestinal cells (Fig. 2A). Together, these results demonstrate that disruption of specific mitochondrial ETC complexes is sufficient to induce the internalization of the heterochromatin reporter in the intestine, recapitulating the regional pattern of reporter positioning observed following *mrg-1* loss.

Mitochondrial dysfunction activates the evolutionarily conserved p38 and JNK mitogen-activated protein kinase (MAPK) stress signaling pathways, which regulate the expression of innate immune genes (Munkácsy et al. 2016; Raingeaud et al. 1995), a gene category that is among the most significantly enriched following *mrg-1* loss (Fig. 1A). We therefore asked whether activation of these signaling pathways alone is sufficient to induce reporter detachment. To this end, we depleted *rpm-1*, which negatively regulates both pathways by targeting the MAPKKKs DLK-1 and MLK-1 (Nix et al. 2011), and *vhp-1*, a MAPK phosphatase that also acts as a negative regulator of these signaling cascades (Munkácsy et al. 2016; Nix et al. 2011). Knockdown of either negative regulator was sufficient to delocalize the reporter from the nuclear periphery in anterior intestinal cells (Fig. 2B). These findings indicate that activation of stress signaling, even in the absence of direct mitochondrial perturbation, is sufficient to alter localization of the reporter and phenocopy the effect of *mrg-1* loss. In contrast to pathway activation, knockdown of individual MAP kinases failed to rescue reporter internalization in *mrg-1*; *cec-4* mutants (Supplemental Fig. S2B), suggesting functional redundancy within and between the p38 and JNK signaling pathways.

Because CBP-3 is required for reporter detachment following loss of MRG-1 (Fig. 1D-1G), we asked whether it is also required when this is induced by mitochondrial dysfunction or activation of stress signaling. We therefore quantified the positioning of the heterochromatin reporter under these conditions in animals carrying either a wildtype (WT) or null *cbp-3* allele. Loss of *cbp-3* fully suppressed reporter internalization induced by depletion of ETC complex III, IV and V components, and the same result was observed following *rpm-1* knockdown (Fig. 2C). Importantly, loss of *cbp-3* had no effect on reporter positioning in *cec-4* single mutants under control RNAi conditions (Fig. 2C), indicating that CBP-3 regulates heterochromatin positioning specifically in response to stress rather than under basal conditions.

Next, to determine whether mitochondrial dysfunction and stress signaling act through the same pathway as *mrg-1* to regulate positioning of the heterochromatic reporter, we performed an epistasis analysis. We depleted one positive hit from each delocalizing ETC complex (*cyc-1*, *cco-4*, and *atp-3*) or *rpm-1* in *mrg-1; cec-4* mutants, in which the reporter is already detached from the nuclear periphery, and quantified reporter positioning. None of these perturbations induced further reporter internalization relative to control RNAi-treated *mrg-1; cec-4* animals (Fig. 2D), consistent with mitochondrial dysfunction and stress signaling acting within the same pathway as loss of *mrg-1* to regulate heterochromatin positioning.

Finally, to exclude the possibility that mitochondrial perturbation affects chromatin organization by reducing MRG-1 abundance, we quantified the protein levels of a functional, endogenously tagged MRG-1-GFP (Doronio et al. 2022). MRG-1 levels remained unchanged following perturbation of ETC complex III by *cyc-1* or *F45H10.2* RNAi or activation of p38/JNK signaling by *rpm-1* or *vhp-1* RNAi (Supplemental Fig. S2C), indicating that stress-induced reporter internalization is not a consequence of reduced MRG-1 expression.

Together, these findings establish that mitochondrial dysfunction and activation of stress signaling are sufficient to induce heterochromatin reporter detachment through a CBP-3-dependent pathway, thereby phenocopying the effect of MRG-1 loss.

### *cbp-3* ablation partially rescues the transcriptional response to MRG-1 loss

CBP-3 regulates the transcriptional response to mitochondrial stress (Munkácsy et al. 2016). Because *mrg-1* mutants exhibit a transcriptional signature consistent with mitochondrial stress (Fig. 1A) and because CBP-3 contributes to the heterochromatin reorganization induced by loss of MRG-1 (Fig. 1D-G), we asked whether CBP-3 also contributes to the gene expression response triggered by MRG-1 depletion. To test this, we performed RNA-seq on L1 larvae exposed to either *mrg-1* or control RNAi in WT and *cbp-3* mutant backgrounds. Importantly, because CBP-3 is highly abundant in the intestine (Fig. 1J), the effects of *cbp-3* loss on gene expression detected by whole-animal RNA-seq are likely to originate from this tissue, where CBP-3 regulates heterochromatin reporter positioning.

Differential expression analysis identified 153 upregulated and 98 downregulated genes following *mrg-1* RNAi (Fig. 3A). As expected, the upregulated genes were enriched for GO categories associated with innate immunity and defense against bacteria (Supplemental Fig. S3A). For genes differentially expressed following MRG-1 depletion, expression changes were generally attenuated in the *cbp-3* mutant background (Fig. 3B and 3C). To quantify this effect, we classified differentially expressed genes according to the extent of their rescue by *cbp-3* loss (see Methods) (Fig. 3D). This analysis revealed that whereas 49% of the genes upregulated upon *mrg-1* depletion were unaffected by loss of *cbp-3* (Fig. 3E), the expression of 35% was at least partially restored in the *cbp-3* mutant background (Fig. 3D and 3E), indicating that mitochondrial stress signaling contributes to a substantial fraction of the transcriptional response induced by MRG-1 loss. Rescue of genes downregulated following *mrg-1* depletion was also observed, with 29% showing at least partial restoration in the absence of *cbp-3* (Fig. 3E).

**Fig. 3.**
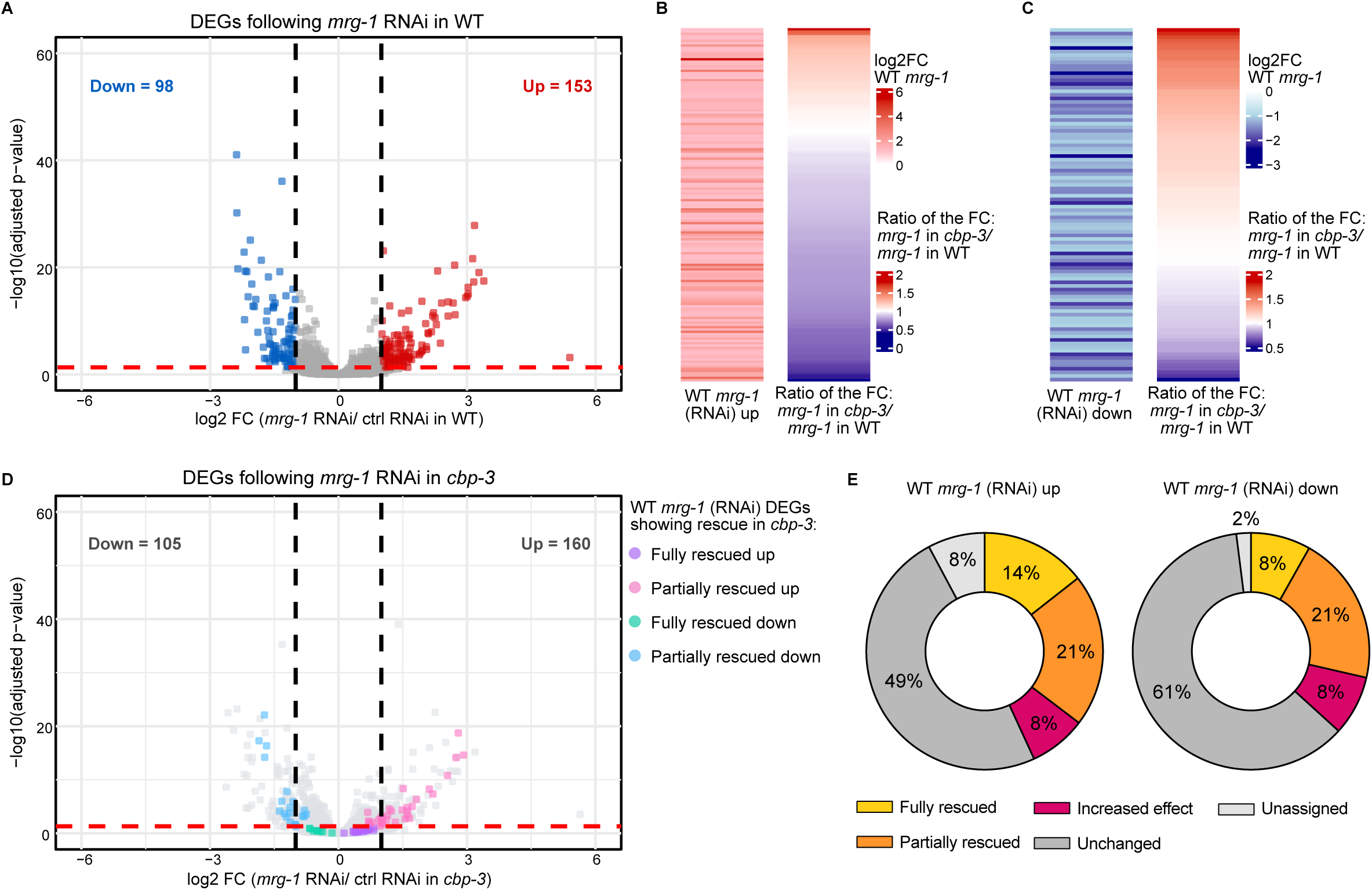
*cbp-3* ablation partially rescues the transcriptional response to MRG-1 loss. **A)** Volcano plot showing genes differentially expressed following *mrg-1* RNAi relative to control RNAi in WT L1 larvae. **B)** Heatmap of the log2 fold change (FC) of genes upregulated following *mrg-1* RNAi in WT relative to control RNAi (left), and the ratio between the FC following *mrg-1* RNAi in *cbp-3* mutants and the FC following *mrg-1* RNAi in WT (right). **C)** Same as (B) but for genes downregulated following *mrg-1* RNAi in WT. **D)** Volcano plot showing gene-expression changes following *mrg-1* RNAi relative to control RNAi in *cbp-3* mutants. Colored points indicate genes identified as differentially expressed following *mrg-1* RNAi in WT in (A) that were classified as fully or partially rescued in *cbp-3* mutants (see Methods for classification criteria). **E)** Donut charts showing the classification of genes differentially expressed following *mrg-1* RNAi in WT in (A) according to their response to *cbp-3* mutation. Categories include fully rescued, partially rescued, increased effect, unchanged, and unassigned.

While loss of *cbp-3* rescued a substantial subset of *mrg-1*-induced differentially expressed genes (DEGs), it also further exacerbated the expression changes of a small fraction of DEGs identified following *mrg-1* depletion in WT animals (Fig. 3E and Supplemental Fig. S3B). Furthermore, 53 upregulated and 17 downregulated genes were differentially expressed exclusively when *mrg-1* was depleted in the *cbp-3* mutant background (Supplemental Fig. S3B). Notably, however, only 24 genes were differentially expressed in *cbp-3* mutants compared to WT animals under control RNAi conditions (Supplemental Fig. S3C), of which only 3 were also differentially expressed following *mrg-1* depletion in WT animals. These findings indicate that the role of CBP-3 in modulating the transcriptional response to *mrg-1* depletion is not simply a consequence of its regulation of the affected genes under basal conditions. Rather, they are consistent with CBP-3 functioning primarily under perturbed cellular conditions, supporting its role as a mediator of stress-induced rather than basal transcriptional responses.

MRG-1 has also been implicated in heterochromatin silencing (Towbin et al. 2012). We therefore asked whether the CBP-3-dependent stress response also contributes to repeat derepression induced by loss of MRG-1. RT-qPCR analysis confirmed that *mrg-1* depletion leads to derepression of specific repetitive elements (Supplemental Fig. S3D). However, repeat silencing was not restored by *cbp-3* mutation (Supplemental Fig. S3D).

Together, these results indicate that CBP-3 contributes to the stress-dependent transcriptional response and heterochromatin reporter detachment induced by loss of MRG-1, whereas repeat silencing is regulated by MRG-1 through a distinct, CBP-3-independent mechanism.

### CBP-3–dependent rescue of nuclear phenotypes in *mrg-1* mutants is associated with detrimental organismal effects

Our results show that loss of *cbp-3* partially rescues both the transcriptional and chromatin organization defects caused by MRG-1 depletion. We next investigated the physiological consequences of *cbp-3* loss in the context of MRG-1 depletion.

Because CBP-3 mediates mitochondrial stress responses (Munkácsy et al. 2016), we assessed mitochondrial function by measuring mitochondrial oxygen consumption rate (OCR) using Seahorse assays (Koopman et al. 2016). Basal mitochondrial OCR was reduced following *mrg-1* depletion in WT animals (Fig. 4A), and although this reduction only approached statistical significance in WT, it became more pronounced and significant in the absence of *cbp-3* (Fig. 4B), while no significant changes in maximal OCR were detected across these conditions (Supplemental Fig. S4A-B). These results are consistent with CBP-3 promoting adaptive responses that mitigate the mitochondrial dysfunction induced by loss of MRG-1.

**Fig. 4.**
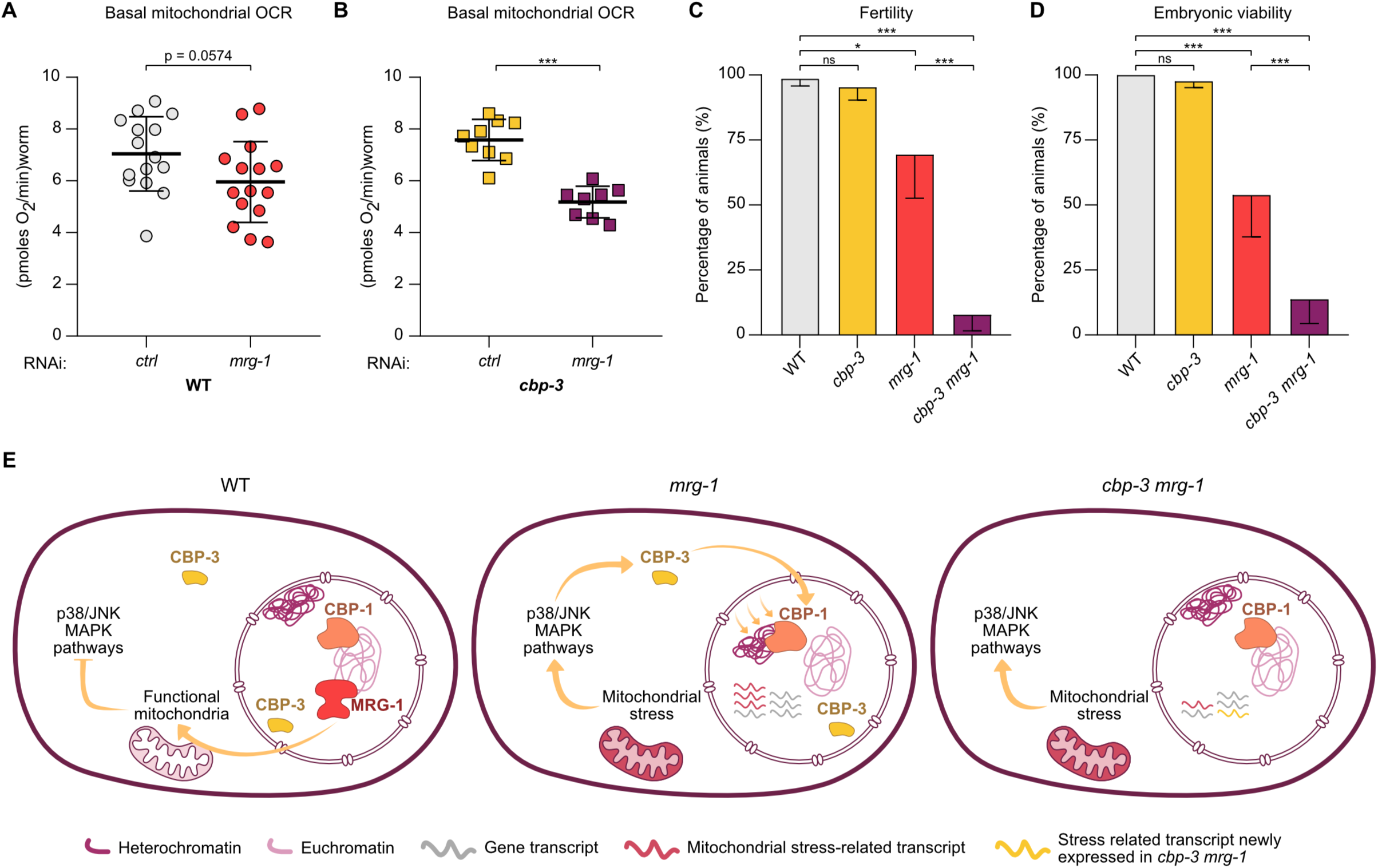
I CBP-3-dependent signaling promotes organismal adaptation to mitochondrial dysfunction following loss of MRG-1. **A)** Basal mitochondrial oxygen consumption rate (OCR) of WT adult worms exposed to mrg-1 RNAi, measured using the Seahorse assay. Individual data points represent technical replicates from 5 independent biological replicates, 20-30 worms per technical replicate. **B)** Same as (A) but in cbp-3 mutants. Results are from 3 biological replicates, 20-30 worms per technical replicate. **C)** Quantification of the percentage of fertile adult worms of the indicated genotypes. For mrg-1 single and cbp-3 mrg-1 double mutants, the first generation of homozygous animals-derived from balanced heterozygous parents-were scored. Results are from 3 biological replicates, n (worms) ≤ 60. Error bars represent standard deviation. **D)** Quantification of the percentage of viable embryos produced by adult worms of the indicated genotypes. For mrg-1 single and cbp-3 mrg-1 double mutants, embryos corresponding to the second generation, produced by first-generation homozygous animals were scored. Results are from 4 biological replicates, n (embryos) ≤ 99. Error bars represent standard deviation. **E)** Model, loss of MRG-1 activates mitochondrial stress signaling, leading to CBP-3-dependent transcriptional and chromatin changes that contribute to organismal adaptation to mitochondrial stress. For A and B, statistical significance was assessed using an unpaired t-test between the two conditions. For C and D, statistical significance was assessed using a one-way ANOVA followed by Sidak’s test for multiple comparisons. *p< 0.05, ***p < 0.001, ns = not significant. Exact p-values and n values are listed in Supplemental Table ST3.

We next investigated how disrupting the CBP-3-dependent stress response affects organismal fitness in *mrg-1* mutants. Specifically, we quantified fertility and embryonic viability in *cbp-3 mrg-1* double mutants and compared them with the corresponding single mutants and WT animals. As *mrg-1* mutants exhibit a maternal-effect sterility phenotype (Fujita et al. 2002), we measured fertility in the first generation of homozygous mutants and embryonic viability in their progeny. While *cbp-3* single mutants display no significant defect compared to WT, and *mrg-1* mutants show a moderate reduction in fertility and viability, *cbp-3 mrg-1* double mutants are nearly completely sterile (Fig. 4C) and produce almost no viable progeny due to severe embryonic lethality (Fig. 4D). As observed for the transcriptional response to *mrg-1* knockdown, the contribution of CBP-3 to these phenotypes became apparent only in the absence of MRG-1, supporting its role in promoting adaptive responses to MRG-1 loss.

Together, these findings indicate that, although loss of CBP-3 partially restores the nuclear phenotypes caused by MRG-1 depletion, it also exacerbates mitochondrial dysfunction and compromises organismal fitness. These observations indicate that the CBP-3-dependent nuclear changes induced by MRG-1 loss contribute to adaptation to mitochondrial stress.

## Discussion

MRG-1 was the first component of the active chromatin compartment shown to regulate heterochromatin positioning (Cabianca et al. 2019). Nonetheless, MRG-1 itself is largely excluded from heterochromatic domains, raising the question of how it indirectly regulates heterochromatin positioning. Here, we identify mitochondrial stress signaling as a previously unrecognized mediator of MRG-1-dependent nuclear changes.

We show that loss of MRG-1 activates a mitochondrial stress response that contributes to heterochromatic reporter detachment (Fig. 1A-G) and to the accompanying gene expression changes (Fig. 3B-E). Furthermore, preventing this stress response partially restores these nuclear phenotypes but compromises mitochondrial function and organismal fitness (Fig. 4A-D), suggesting that the changes in genome regulation elicited by MRG-1 loss are, at least in part, components of an adaptive response to mitochondrial stress.

A central question raised by our findings is how loss of a chromatin reader activates mitochondrial stress signaling. Although mammalian MRG15 has been reported to localize to mitochondria in the liver (Tian et al. 2022), we found no evidence for mitochondrial localization of a functional endogenously tagged MRG-1 in the *C. elegans* intestine (data not shown). While we cannot exclude mitochondrial localization below the detection limit of confocal microscopy, we favor an indirect mechanism. Given the established role of MRG-1 in chromatin complexes, mitochondrial stress may arise through altered expression of genes involved in mitochondrial homeostasis, metabolism, or stress responses, that are regulated by MRG-1-containing complexes. Consistent with this possibility, loss of the transcriptional corepressor SIN-3 – which forms a conserved complex with MRG-1 (Beurton et al. 2019; Wan et al. 2023) – directly deregulates the expression of numerous mitochondrial genes and results in mitochondrial dysfunction (Giovannetti et al. 2024), raising the possibility that MRG-1 contributes to mitochondrial homeostasis through its nuclear role within the SIN3 complex.

Basal respiration is only mildly reduced following MRG-1 depletion, but when the CBP-3-dependent signaling pathway is impaired in *mrg-1* mutants, this exacerbates mitochondrial dysfunction, fertility defects and embryonic lethality. These observations suggest that part of the transcriptional and chromatin changes induced by MRG-1 loss contribute to adaptation to mitochondrial stress.

The exact nature of the mitochondrial alteration induced by loss of MRG-1 remains to be determined. However, the observation that only perturbation of ETC complexes III, IV and V – but not complexes I or II – induces delocalization of the heterochromatic reporter may provide an important clue. Whereas complexes I and II provide alternative entry points into the electron transport chain, complexes III, IV and V constitute obligatory downstream steps of oxidative phosphorylation (Vercellino and Sazanov 2022). This raises the possibility that a more complete impairment of electron transport, or downstream consequences associated with ETC dysfunction, may be required to trigger 3D chromatin reorganization. At the same time, because activation of p38 signaling alone is sufficient to detach the heterochromatic reporter from the nuclear periphery, other cellular stresses that activate this pathway may similarly influence heterochromatin positioning.

Although ATFS-1 regulates more than half of the transcriptional response to mitochondrial stress (Nargund et al. 2012), *atfs-1* depletion did not restore heterochromatin reporter localization at the nuclear periphery, whereas *cbp-3* knockdown did (Fig. 1D–G). *cbp-3* was long considered a pseudogene but has recently been shown to regulate promoter activity (Horowitz et al. 2023) and to be required for an ATFS-1-independent PMK-3/MAPK mitochondria-to-nucleus signaling pathway (Munkácsy et al. 2016). Notably, both studies relied exclusively on RNAi-mediated depletion of *cbp-3*. Here, we provide the first genetic evidence that *cbp-3* is a functional gene by using a mutant allele. While CBP-3 is largely dispensable for heterochromatin positioning, gene expression, and organismal fitness under normal growth conditions, it promotes heterochromatic reporter detachment in *mrg-1; cec-4* double mutants, mediates approximately one-third of the transcriptional changes induced by loss of MRG-1, and is required to maintain fertility and embryonic viability in the absence of MRG-1. The selective requirement for CBP-3 under conditions of MRG-1 loss is consistent with its established role in stress-responsive signaling pathways (Munkácsy et al. 2016).

While our data strongly support a role for mitochondrial stress in regulating the nuclear changes induced by loss of MRG-1, they also indicate that this pathway explains only part of the consequences of MRG-1 depletion. Although loss of *cbp-3* completely restores reporter perinuclear binding following ETC perturbation, it only partially rescues reporter positioning and gene expression changes following MRG-1 depletion. Moreover, repeats de-repression caused by MRG-1 depletion persists in the absence of CBP-3. Together, these findings indicate that MRG-1 regulates distinct aspects of genome function through different downstream pathways. Whether the remaining effects arise from direct chromatin-associated functions of MRG-1 or from additional branches of the cellular stress response remains an important question for future studies.

It was previously shown in *C. elegans* that mitochondrial stress induces chromatin changes through the heterochromatic H3K9 methyl transferase MET-2 (Tian et al. 2016), consistent with heterochromatin being remodeled in response to mitochondrial dysfunction. Interestingly, dysfunctional mitochondrial quality control leads to ETC defects that engage a retrograde (mitochondria-to-nucleus) signaling program, inducing chromatin remodeling, cell dedifferentiation and impairing the function of metabolic tissues (Walker et al. 2025). As heterochromatin stabilizes cell differentiation programs (Burton and Torres-Padilla 2021) and its accumulation at the nuclear periphery accompanies cell differentiation (Leung et al. 1999; Rozman et al. 1993; Solovei et al. 2013; Ugarte et al. 2015), an intriguing possibility is that mitochondrial stress-induced heterochromatin reorganization may contribute to changes in cell identity.

In sum, our work identifies mitochondrial stress signaling as a previously unrecognized mediator of the nuclear changes induced by loss of the chromatin reader MRG-1. Although the impact of mitochondrial stress on endogenous genome–nuclear lamina interactions remain to be determined, our findings support the idea that cellular stress responses can shape nuclear organization.

## Material and Methods

### C. elegans maintenance and strains

*C. elegans* nematodes were maintained at 20 °C on Nematode Growth Media (NGM) agar plates seeded with *Escherichia coli* OP50 bacteria culture, except for RNAi experiments (See RNAi treatments). All *C. elegans* strains used are listed in Supplemental Table ST1.

### RNAi treatments

RNAi experiments were performed at 20 °C on NGM plates supplemented with 100 mM carbenicillin (6344.2, Roth) and 100 µM isopropyl-β-D-thiogalactoside (IPTG; A4773,0005, AppliChem) seeded with double-stranded RNA-producing bacteria, grown overnight at 37 °C. As negative control, the RNAi clone containing the empty L4440 vector (Fire vector library) was used. In every experiment, *let-607* RNAi was performed in parallel to confirm the efficiency of the RNAi plates, as a phenotype of very delayed growth/lethality is expected compared with worms grown on L4440 RNAi. Except for *atfs-1* (kind gift of the Auxwerx lab, EPFL Lausanne, Switzerland), all RNAi clones used were obtained from the Ahringer library (Source BioScience Ltd.). All RNAi clones used were sequenced to confirm target specificity. Every experiment was performed at least two times using biologically independent samples and RNAi bacteria cultures.

For *C. elegans* strains without balancer chromosome, embryos or synchronized L1s obtained by hypochlorite treatment were seeded on RNAi plates and grown for 3.5 to 4 days, then adults and larvae were washed off the plate to leave only unhatched embryos. 2-3 hours later, newly hatched L1s were collected with M9 for imaging. For RNAi-mediated depletion of ETC complex IV and V components, worms were initially maintained on L4440 control RNAi and transferred at the L4 stage to plates seeded with the corresponding RNAi bacteria. This strategy prevented early developmental arrest, allowing the animals to produce progeny for imaging.

For experiments involving strains maintained with a chromosome balancer (i.e., strains carrying an *mrg-1* mutant allele), gravid balanced heterozygous adults were manually transferred to RNAi plates. Approximately 72 hours later, homozygous mutant gravid adults lacking the balancer chromosome (identified by their WT-like locomotion) were manually transferred to fresh RNAi plates. The following day, the adults were removed by washing, leaving only unhatched embryos. Newly hatched L1 larvae were collected 2–3 hours after for imaging.

For RNAi-mediated depletion of ETC complex IV and V components in strains carrying a balancer chromosome (*mrg-1* KO), heterozygous adults with the balancer were initially maintained on L4440 control RNAi. Homozygous mutant gravid adults were then transferred to the corresponding RNAi plates approximately 30 hours before imaging. L1 larvae were collected for imaging as described above.

### Microscopy

Microscopy was performed using a live-cell imaging confocal spinning-disk microscope from Visitron Systems GmbH, equipped with the following instruments: Nikon Eclipse Ti2 microscope with a Plan apo λ 100x/1,45 oil objective, a Plan apo λ 60x/1,45 oil objective, a Plan Apo λ 10× objective, VS-Homogenizer, Electron multiplying CCD camera (Andor iXon Series) and VisiView software for acquisition.

For image acquisition, freshly hatched L1 larvae were collected in M9 buffer and mounted on 2% agarose pads containing 0.15% sodium azide (Interchim, NJK63A) to immobilize the animals. Adults were hand-picked into an M9 drop directly on the agarose pads. Images used for heterochromatin reporter–nuclear envelope distance measurements and fluorescence intensity analyses were acquired using a Plan Apo λ 100×/1.45 oil immersion objective. Image stacks consisted of 60–75 optical sections collected at 200 nm intervals for the reporter-related analysis, and exactly 65 z-stacks for all images used for MRG-1-GFP fluorescence intensity quantification. For Fig. 1J, adults were imaged at a single focal plane with a Plan Apo λ 10× objective, and L1s with a Plan apo λ 60×/1.40 oil objective.

### Image analysis

Heterochromatin reporter (*gwIs4*) reporter distribution was assessed in different intestinal compartments of L1 larvae using Fiji/ImageJ and the PointPicker plugin (http://bigwww.epfl.ch/thevenaz/pointpicker/), as previously described (Meister et al. 2010a). The position of the GFP-tagged heterochromatin reporter was determined by measuring its shortest distance to the fluorescently labeled inner nuclear membrane and normalizing this value to the nuclear radius in the plane of focus. The uppermost and lowermost focal sections of each nucleus were excluded because of their poor resolution. For visualization, the outermost one-third of the nucleus is highlighted in grey in the plots showing reporter relative radial distribution. This facilitates comparison with the zoning assay used in previous studies (Cabianca et al. 2019), which classifies reporter localization into three nuclear zones of equal area, with reporter detachment corresponding to displacement beyond the outermost peripheral zone.

MRG-1-GFP fluorescence intensity quantification was performed by projecting the mean of all 65 z-stacks, then the mid-body of the worm (corresponding to the portion where the intestine is located) was segmented and fluorescence intensity measured only from this fragment. This was done individually for each worm.

### RNA samples collection and extraction

For each condition, approximately 9,000 synchronized N2 or *cbp-3* mutant L1 larvae, obtained by hypochlorite treatment, were seeded onto 9×10 cm RNAi plates (1000 L1s per plate) containing L4440 control or *mrg-1* RNAi bacteria. After 3.5 days, gravid adults were collected from all conditions and subjected to hypochlorite treatment to isolate embryos, which were allowed to hatch overnight at 20 °C in M9 buffer. Synchronized L1 larvae were then refed for 2.5 hours on plates containing the corresponding RNAi bacteria, collected, washed five times with M9 buffer, resuspended in 500 µL TRIzol reagent (Thermo Fisher Scientific, 15596026), snap-frozen in liquid nitrogen, and stored at −80 °C until RNA extraction.

For RNA extraction, samples were freeze-cracked by five cycles of alternating immersion in liquid nitrogen and a 42 °C water bath, followed by vortexing for 2.5 minutes at room temperature using five 30 seconds on/off cycles. Total RNA was extracted using the RNeasy Mini Kit (74104, Qiagen) according to the manufacturer’s instructions, including a 15 minutes on-column DNase I digestion (79254, Qiagen). RNA concentration was determined using a NanoDrop spectrophotometer, and RNA integrity was assessed using an RNA Pico Chip on a 2100 Bioanalyzer (Agilent Technologies).

### RNA sequencing

For each condition, 1–2 µg of total RNA was diluted in RNase-free water and sent to Novogene GmbH for quality control, library preparation, and paired-end RNA sequencing. Libraries were prepared from 500 ng of input RNA using the TruSeq Stranded Total RNA Library Prep Kit according to the manufacturer’s instructions. This protocol captures messenger RNAs and long non-coding RNAs following ribosomal RNA depletion. Sequencing was performed on a NovaSeq 6000 or NovaSeq X Plus platform using 150 bp paired-end reads.

### RNA sequencing data analysis of differentially expressed genes

RNA-seq data were processed using Bash scripts. Read quality was initially assessed with FastQC, followed by adapter and quality trimming using Trim Galore (https://github.com/FelixKrueger/TrimGalore). Reads were aligned to the *C. elegans* reference genome (ce11) using STAR, retaining multimapping reads (--winAnchorMultimapNmax 200 and --outFilterMultimapNmax 100). Gene-level read counts were generated using featureCounts (Liao et al. 2014), and final quality control was performed with MultiQC (Ewels et al. 2016). Following alignment, only uniquely mapped reads were used for gene quantification and all subsequent analyses. Downstream analyses included only genes with at least 30 reads in a minimum of 3 samples and a pseudocount of 8 was added to minimize large expression changes driven by low read counts, as previously done (Cabianca et al. 2019). Batch effects between biological replicates were corrected using ComBat-seq (Zhang et al. 2020). Differential gene expression analysis was performed using DESeq2 (Love et al. 2014), with WT and *cbp-3* knockout samples analyzed separately to identify genes differentially expressed between control and *mrg-1* RNAi conditions within each genotype. Genes were considered differentially expressed if they exhibited a log2 fold change > 1 (upregulated) or < − 1 (downregulated), with an adjusted *p-*value (padj) < 0.05.

### RNA sequencing data analysis of cbp-3 effect over mrg-1 knock-down

To assess the effect of *cbp-3* loss on the gene expression changes induced by *mrg-1* knockdown, each gene that was differentially expressed following *mrg-1* RNAi in WT animals was classified according to the change in its expression when *mrg-1* RNAi was performed in the *cbp-3* mutant background. Classification was based on the log2 fold change (log2FC), adjusted *P* value (padj), and the fold change (FC) ratio between genotypes (FC(*cbp-3 mrg-1*) / FC(WT *mrg-1*)). All genes were thus sorted between 5 different categories: unchanged, partially rescued, increased effect, fully rescued and unassigned, as outlined below.

We selected differentially expressed genes upon *mrg-1* RNAi in WT (│log2FC │> 1, padj < 0.05). Of these, we considered as “unchanged” (either up or down) genes that remained significant in *cbp-3 mrg-1* RNAi (padj < 0.05) and whose FC difference over WT *mrg-1* RNAi was lower than 20%. We considered as “partially rescued” genes that remained significant in *cbp-3 mrg-1* RNAi (padj < 0.05) and whose FC difference over WT *mrg-1* RNAi was greater than 20%, reducing the effect observed upon *mrg-1* RNAi in WT. We considered as “increased effect” genes that remained significant in *cbp-3 mrg-1* RNAi (padj < 0.05) and whose FC difference over WT *mrg-1* RNAi was greater than 20%, but increasing the effect observed upon *mrg-1* RNAi in WT. To identify “fully rescued” genes, we applied more stringent criteria. First, we considered genes to be “fully rescued” when they were no longer significant in *cbp-3 mrg-1* RNAi, but with a padj > 0.1, thereby applying a more stringent selection. Second, fully rescued genes were defined by a greater than 10 fold ratio of adjusted *P* values between *cbp-3 mrg-1* and WT *mrg-1* (padj(*cbp-3 mrg-1*)/padj(WT *mrg-1*) > 10). Genes defined as “unassigned” had a padj in *cbp-3 mrg-1* between 0.05 and 0.1, or a ratio of adjusted *P* values (padj(*cbp-3 mrg-1*)/padj(WT *mrg-1*) < 10).

### Quantitative PCR with reverse transcription

Complementary DNA (cDNA) was synthesized from either 500 or 1,000 ng of total RNA using the Maxima H Minus cDNA Synthesis Master Mix (M1661, Thermo Fisher Scientific). Gene expression was quantified by real-time PCR using a LightCycler 480 instrument (Roche) and the PowerUp SYBR Green Master Mix (A25742, Thermo Fisher Scientific). Relative expression levels were normalized to the housekeeping gene *cdc-42* by calculating ΔCt values for each sample. Fold changes relative to the control genotype or condition were calculated using the 2-ΔΔCt method. Primer sequences are listed in Supplemental Table ST2.

### Seahorse assay

RNAi bacterial cultures were grown overnight in LB containing tetracycline and ampicillin. The following day, cultures were diluted into LB containing ampicillin only and grown at 37 °C for 3 h. Expression of double-stranded RNA was induced by adding IPTG to a final concentration of 2 mM prior to seeding NGM plates, which were then incubated overnight at room temperature. Worms were subsequently maintained at 20 °C on these plates seeded with HT115(DE3) bacteria carrying either the L4440 empty vector or the *mrg-1* RNAi construct and supplemented with 25 µg/mL carbenicillin and 5 µg/mL nystatin. Oxygen consumption was measured using an Agilent Seahorse XFp Analyzer. Animals were synchronized by allowing gravid adults to lay eggs for 2–3 hours before removing the adults from the plates. Day 5 adult worms (20–30 animals per well) were transferred into Seahorse XFp cell culture miniplates containing M9 buffer. Carbonyl cyanide 4-(trifluoromethoxy) phenylhydrazone (FCCP; Sigma-Aldrich) was prepared in M9 at a final concentration of 250 μM and injected during the assay to force maximal respiration. Sodium azide (NaN₃; Sigma-Aldrich) was prepared in M9 at a final concentration of 400 mM and injected during the assay to inhibit mitochondrial respiration. Oxygen consumption rate (OCR) was measured over eight cycles under basal conditions, followed by ten cycles after FCCP injection and four cycles after NaN₃ injection. Each measurement cycle consisted of 1 minute mixing, 2 minutes waiting, and 3 minutes measurement. Three to five independent biological replicates were performed. OCR values were normalized to the number of worms per well.

### Fertility assessment

For strains lacking a balancer chromosome (N2 and *cbp-3* mutants), 20 day 1 adult worms were individually transferred to NGM plates seeded with OP50. For strains carrying a balancer chromosome (*mrg-1* single mutants and *cbp-3 mrg-1* double mutants), 25 balanced heterozygous adults (Roller phenotype) were first isolated. After 3–4 days, at least 20 first-generation day 1 adult homozygous mutants (WT-like locomotion) were individually transferred to NGM plates seeded with OP50. Fertility was assessed 24 hours after transfer. Animals producing progeny (embryos and/or larvae) were scored as fertile, whereas animals producing no progeny were scored as sterile.

### Embryonic viability assessment

For strains lacking a balancer chromosome (N2 and *cbp-3* mutants), 20-35 embryos laid by day 1 adults were transferred to NGM plates seeded with OP50. For strains carrying a balancer chromosome (*mrg-1* single mutants and *cbp-3 mrg-1* double mutants), balanced heterozygous adults (Roller phenotype) were first isolated. After 3–4 days, first-generation day 1 adult homozygous mutants (WT-like locomotion) were transferred to fresh NGM plates seeded with OP50 and allowed to lay embryos. Few hours later, 20-50 embryos were transferred to NGM plates. The embryos were counted immediately after transfer. After 24 hours, the number of unhatched embryos was recorded, and embryonic viability was calculated as the number of hatched embryos relative to the total number of embryos initially transferred.

### Statistics

For heterochromatin reporter radial distance measurements, MRG-1–GFP fluorescence intensity, Seahorse assays, fertility and embryonic viability assays, data was tested for normality using the Shapiro–Wilk test. Depending on the outcome, either parametric or non-parametric statistical test was applied, associated with a multiple-comparison test when applicable.

The exact statistical tests used for each analysis are indicated in the corresponding figure legends and summarized in Supplemental Table ST3.

## Acknowledgements

We thank the Kelly and Gasser labs for sharing strains and the Auxwerx lab for sharing the *atfs-1* RNAi bacteria clone. Some strains were provided by the CGC, which is funded by NIH Office of Research Infrastructure Programs (P40 OD010440). WormBase (Sternberg et al. 2024) and WormAtlas (supported by NIH OD 010943). The *3xHA-degron-linker-NeonGreen-spacer-cbp-3* expressing strain (strain PHX6880 [*cbp-3* (*syb6880*)]) was obtained by SunyBiotech. This study was funded by the Deutsche Forschungsgemeinschaft (DFG) – Project numbers 459488476 and 507935196 to DSC. MAS contributed with funding from the Spanish Ministerio de Ciencia, Innovación y Universidades, the Spanish Agencia Estatal de Investigación (Project PID2022-139772NB-I00). DSC thanks Helmholtz Munich for support.

## Author contributions

C.Z. and D.S.C conceived the study. C.Z., F.R.P. and MJ.R.P performed the experiments and analyzed the results. C.Z., D.S.C. P.M. and M.A.S. interpreted the results. D.S.C. and M.A.S. provided funding. D.S.C. supervised the study. C.Z. and D.S.C wrote the manuscript. F.R.P. edited the manuscript.

## Supplemental material

**Supplemental Table ST1.**
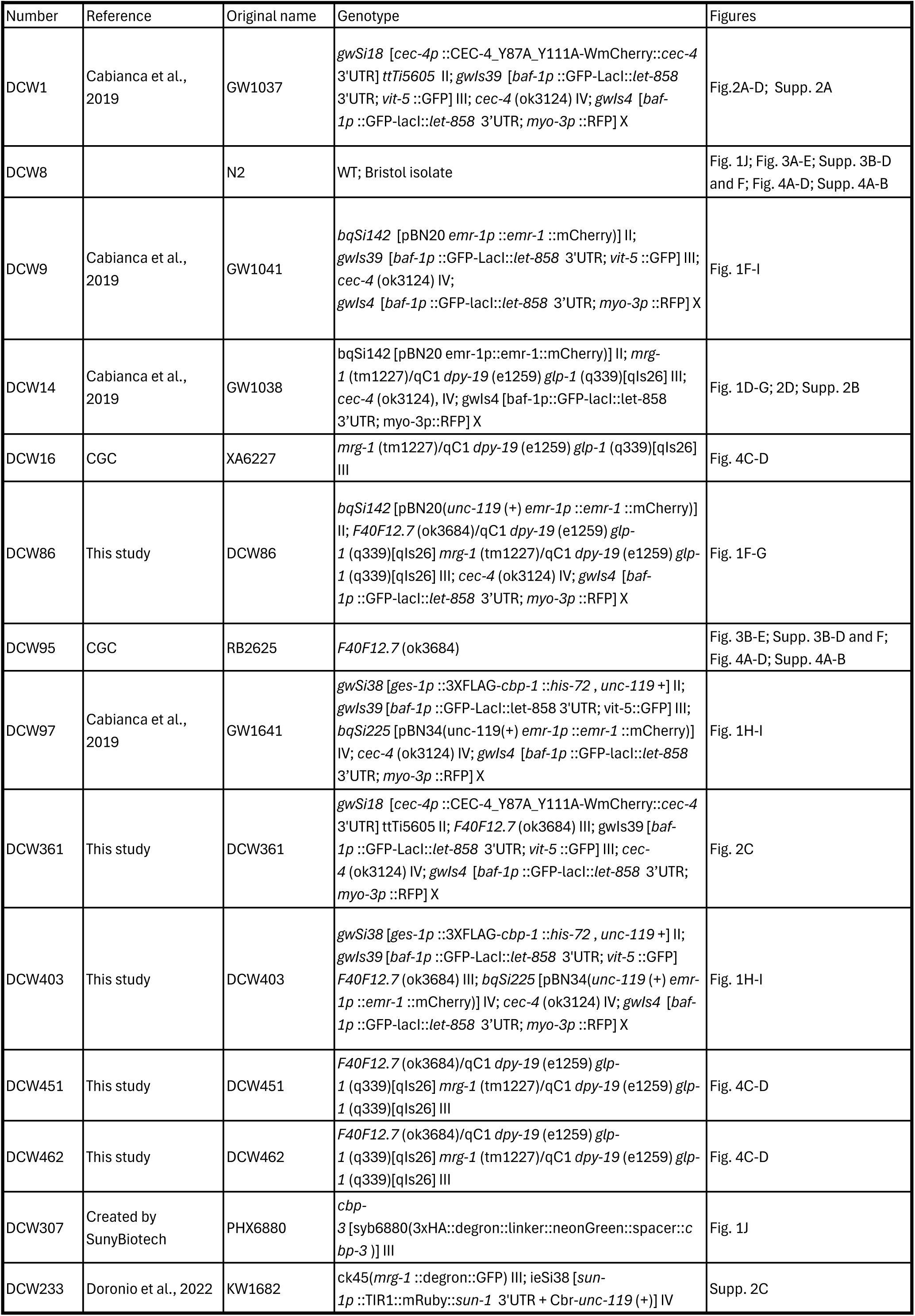
C. elegans strains.

**Supplemental Table ST2.**
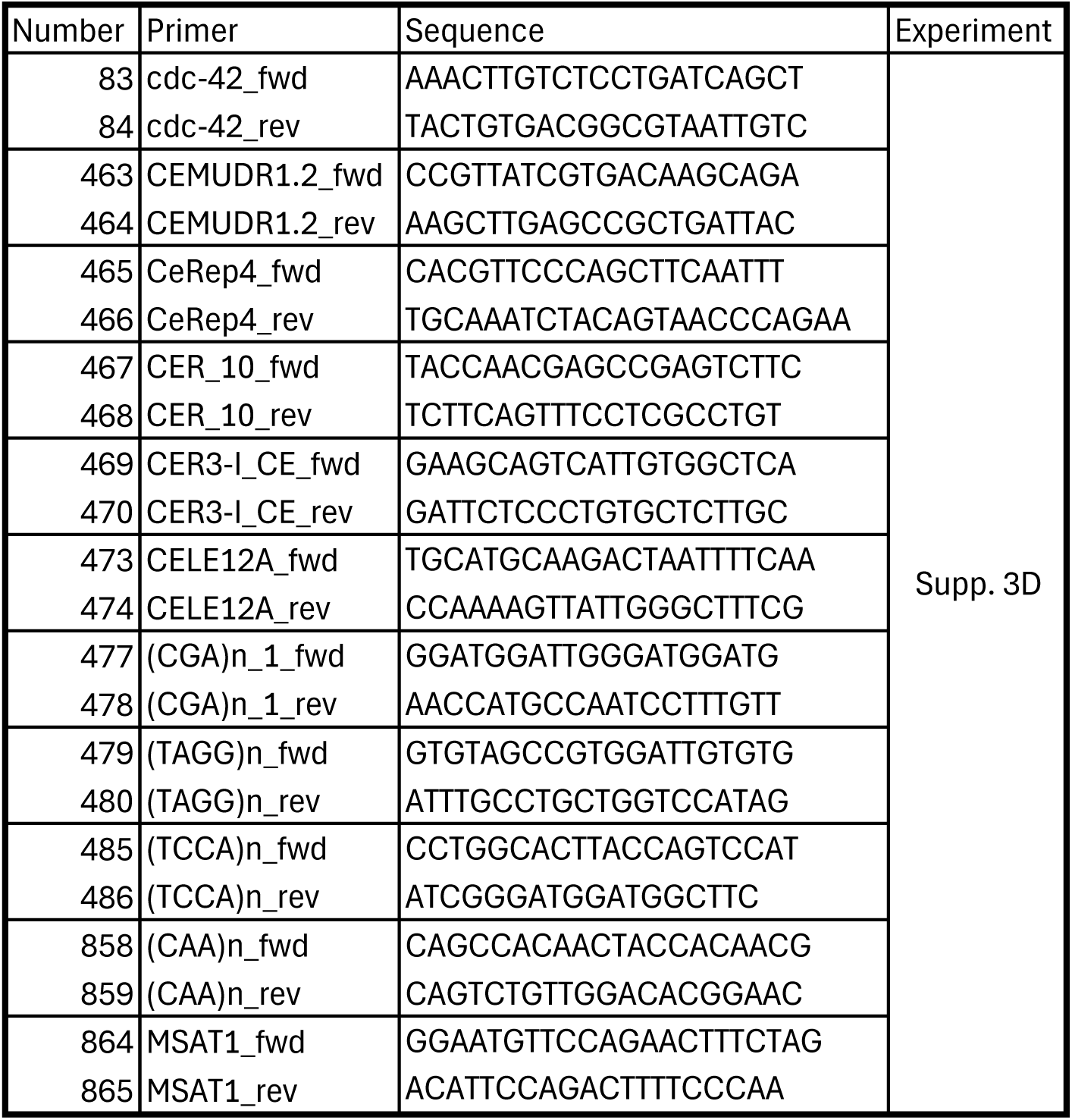
Primers.

**SupplementalTable ST3.**
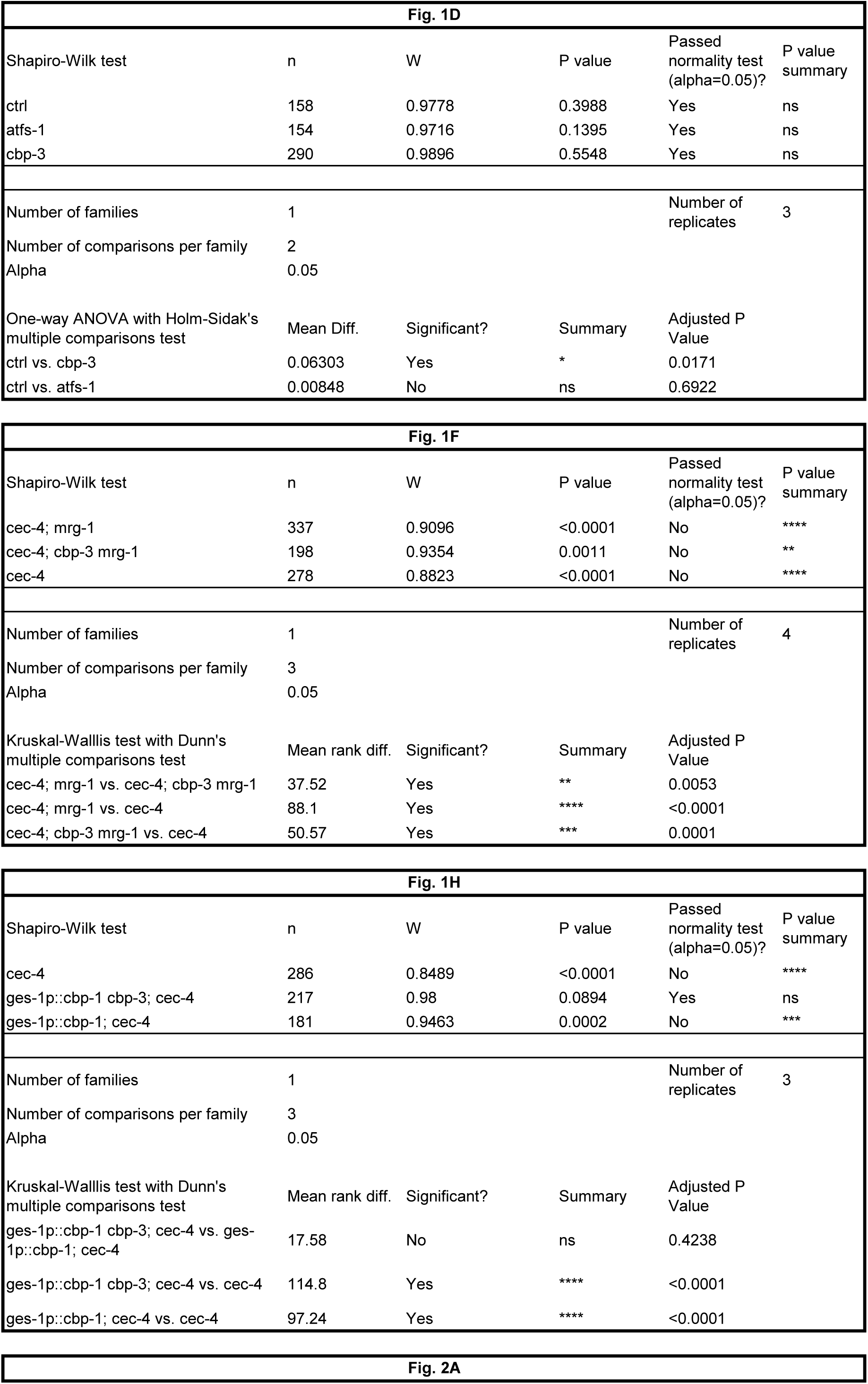
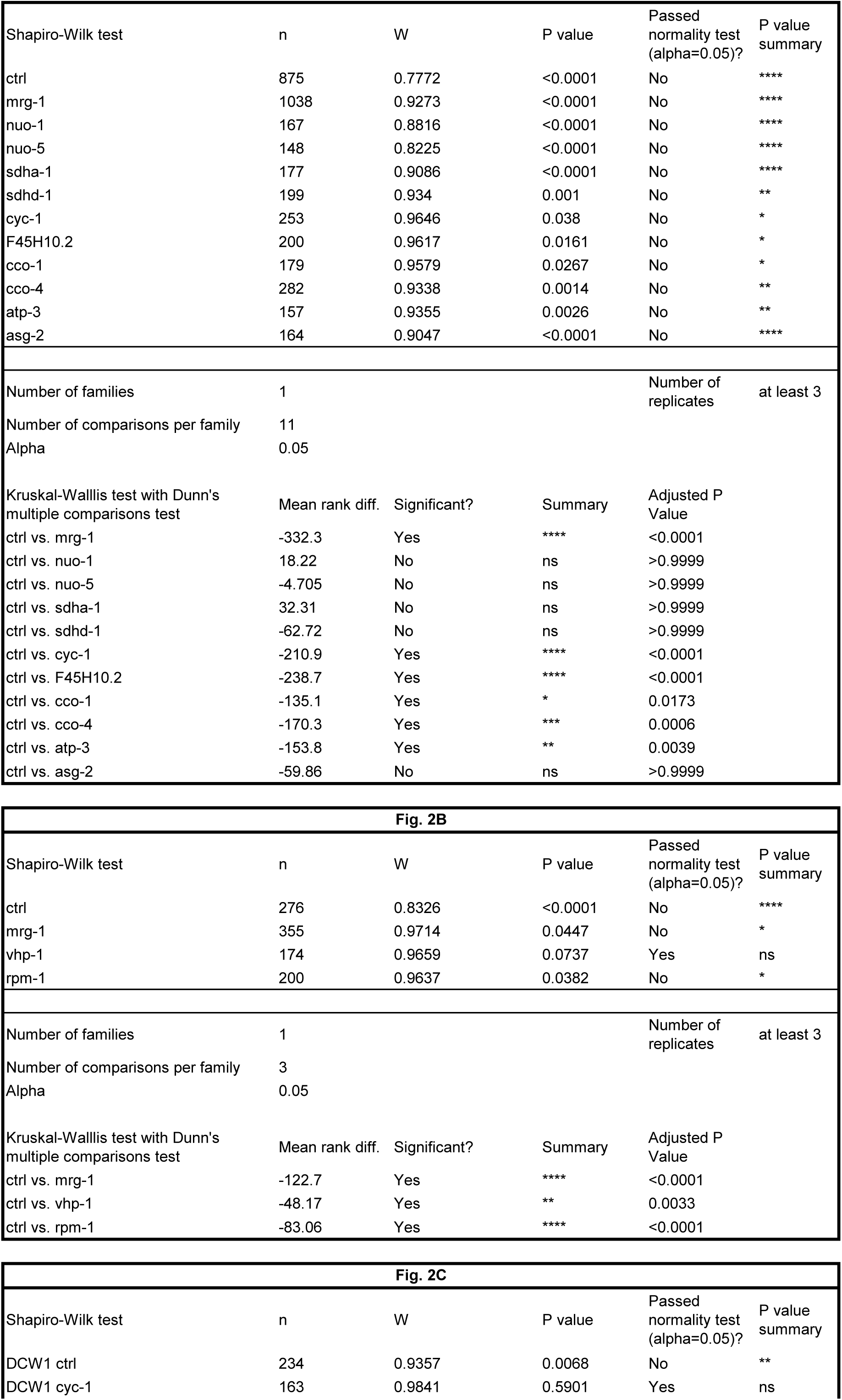
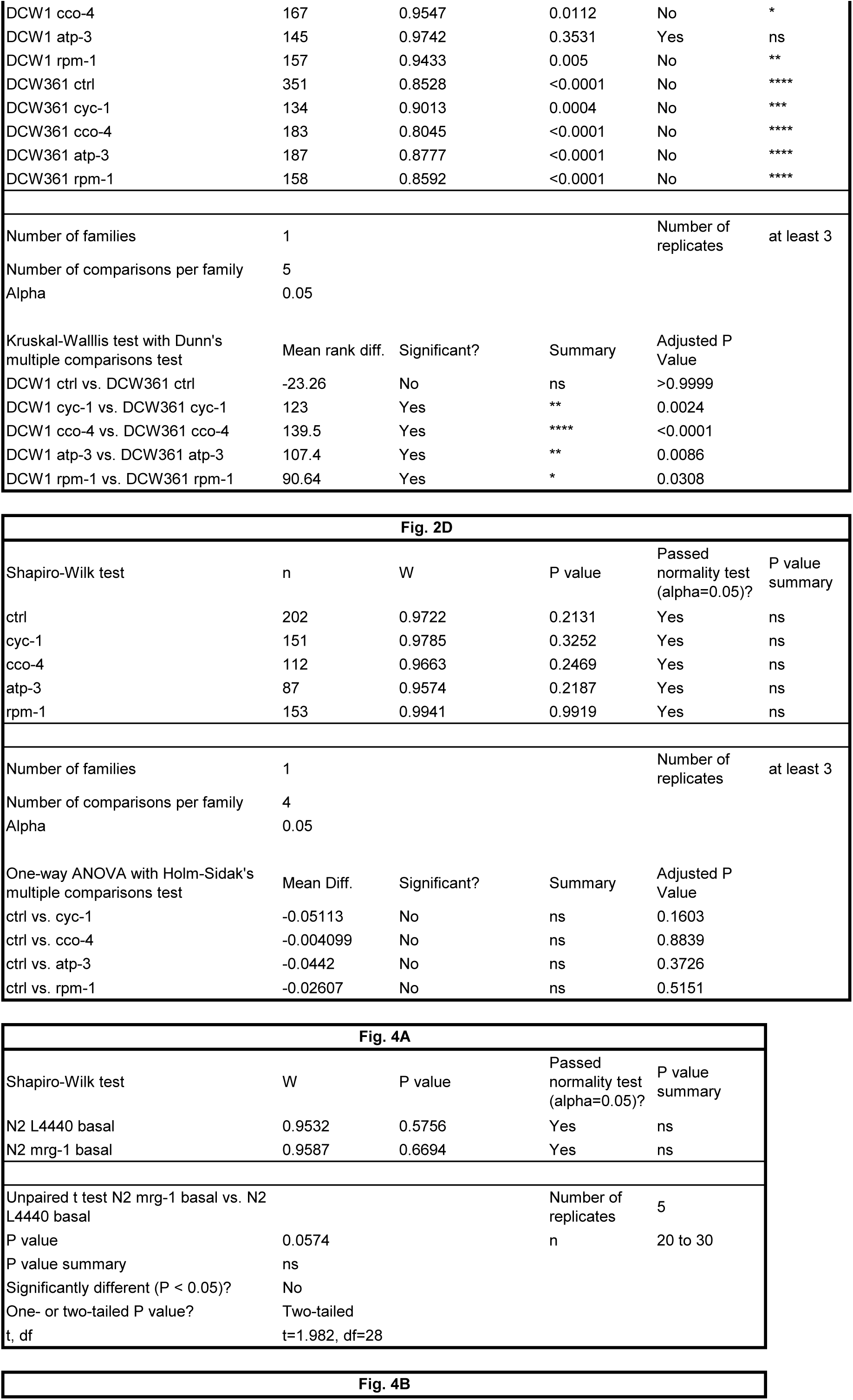
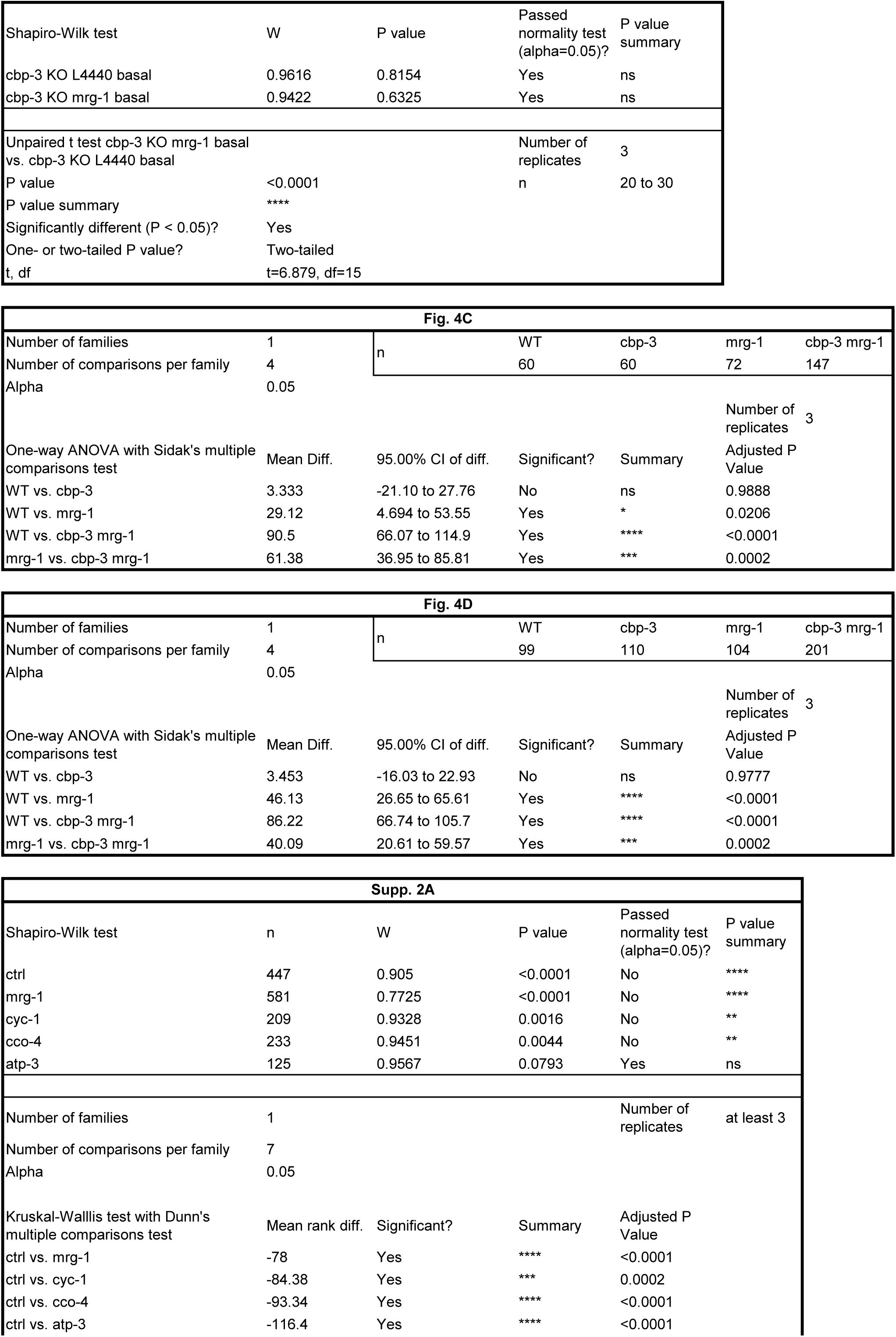
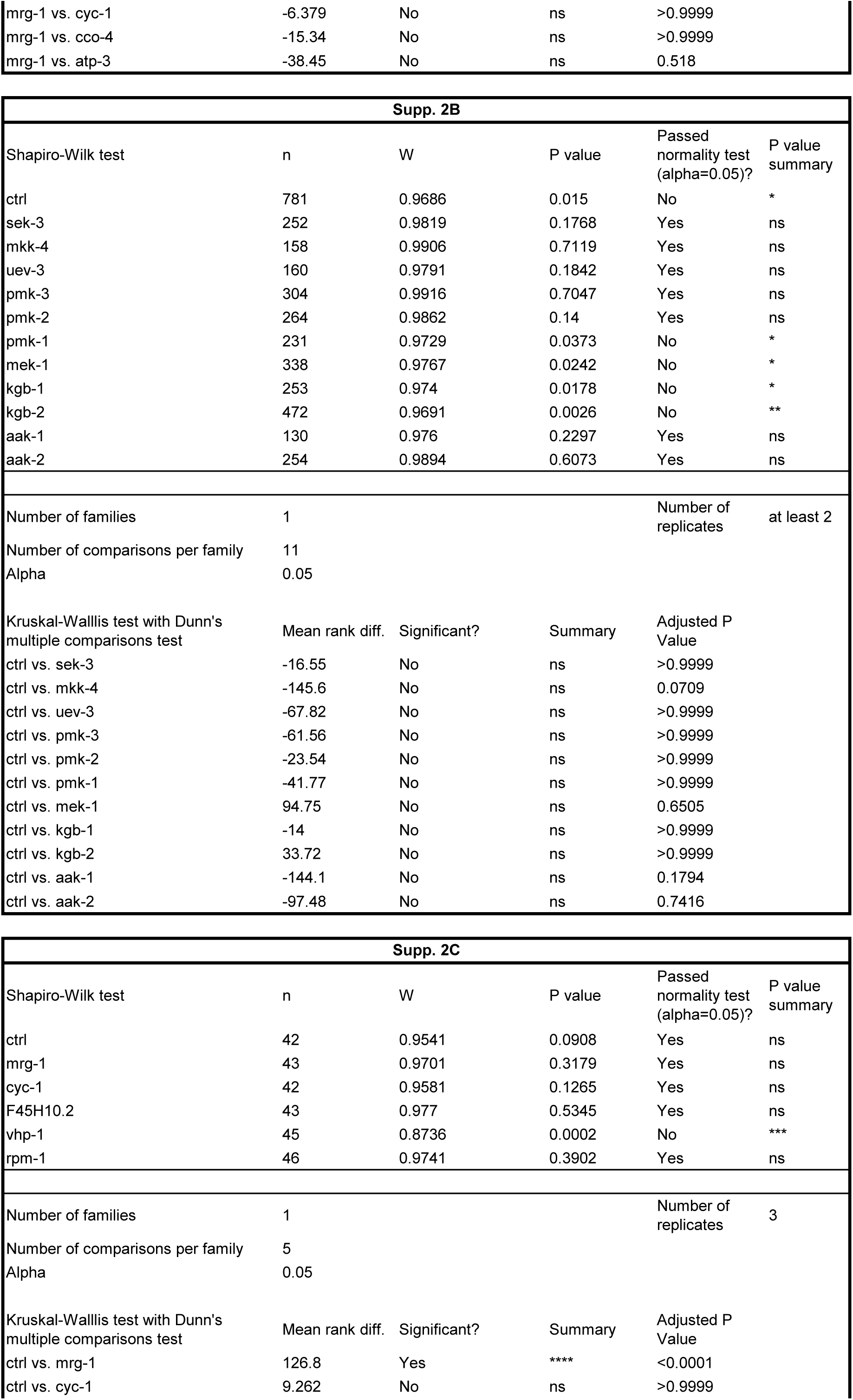
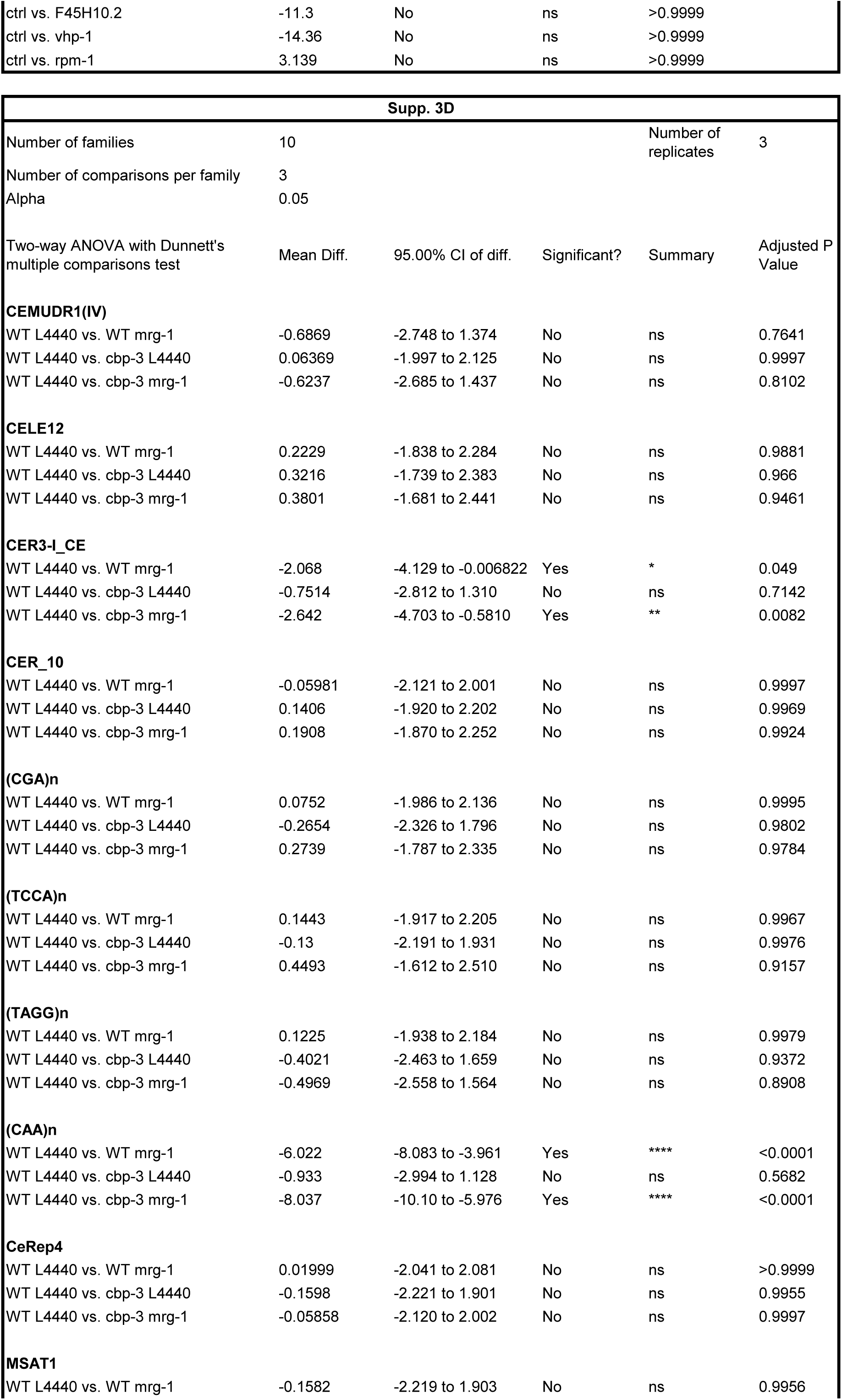
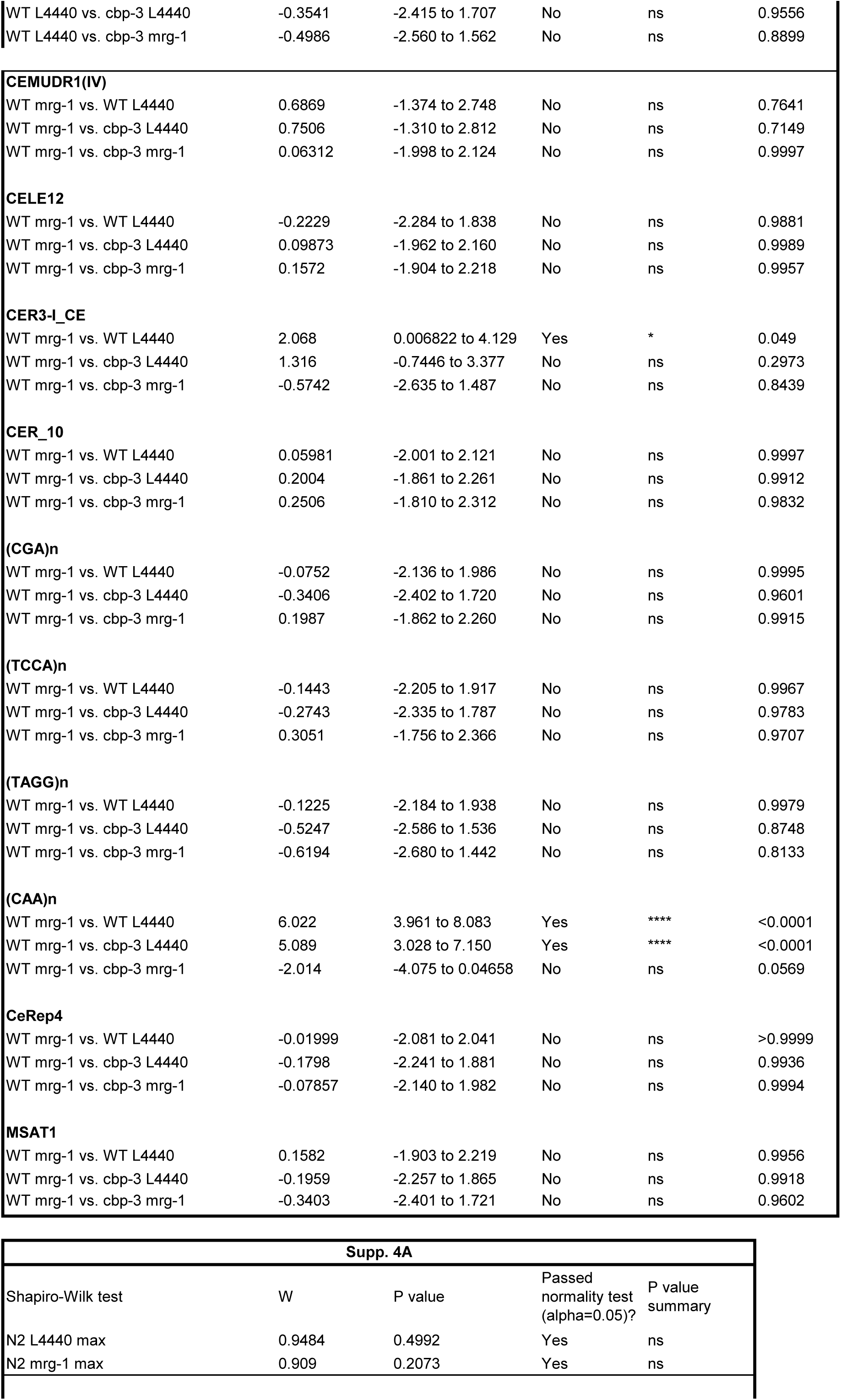
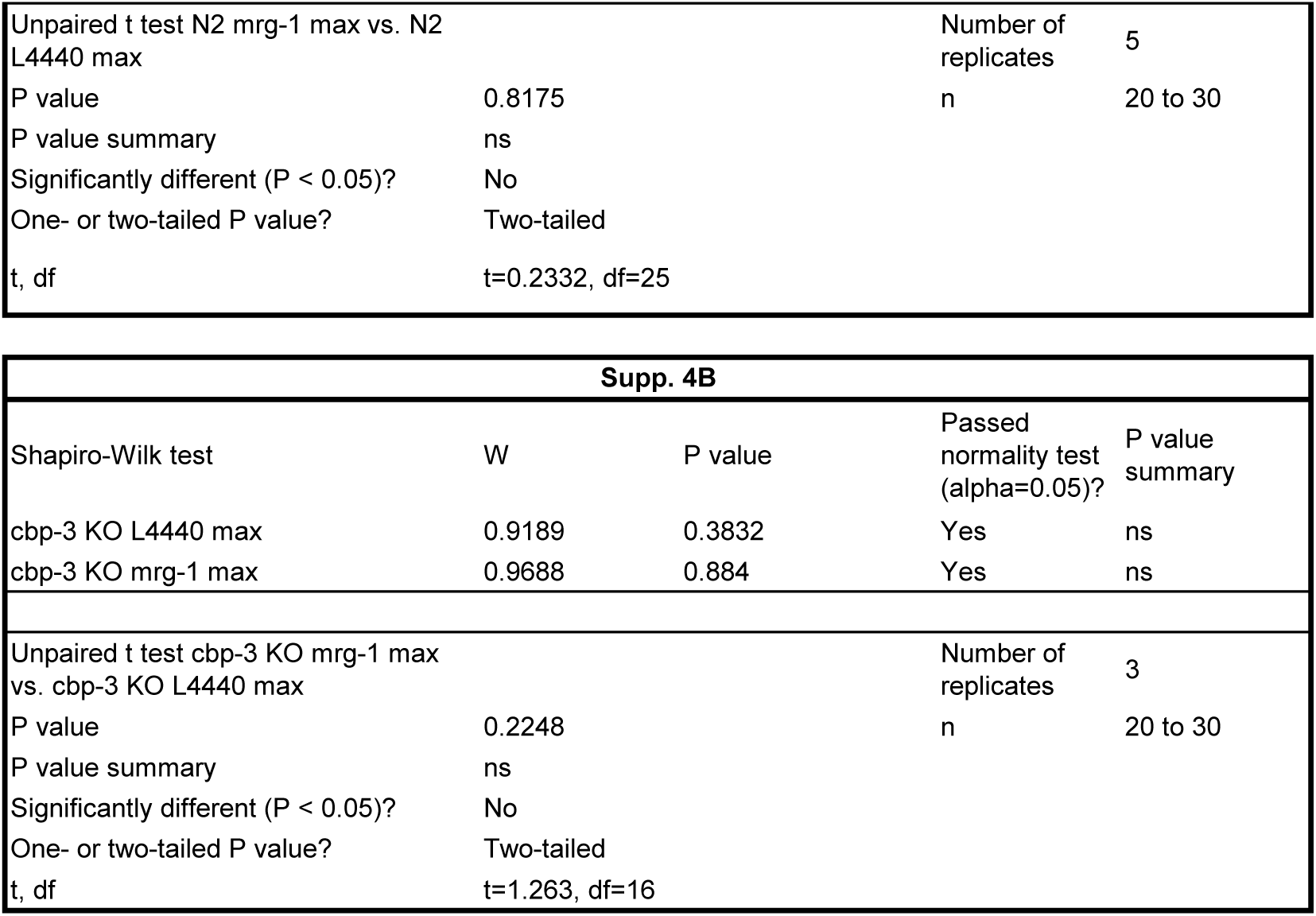
Statistics.

**Supplemental S1.**
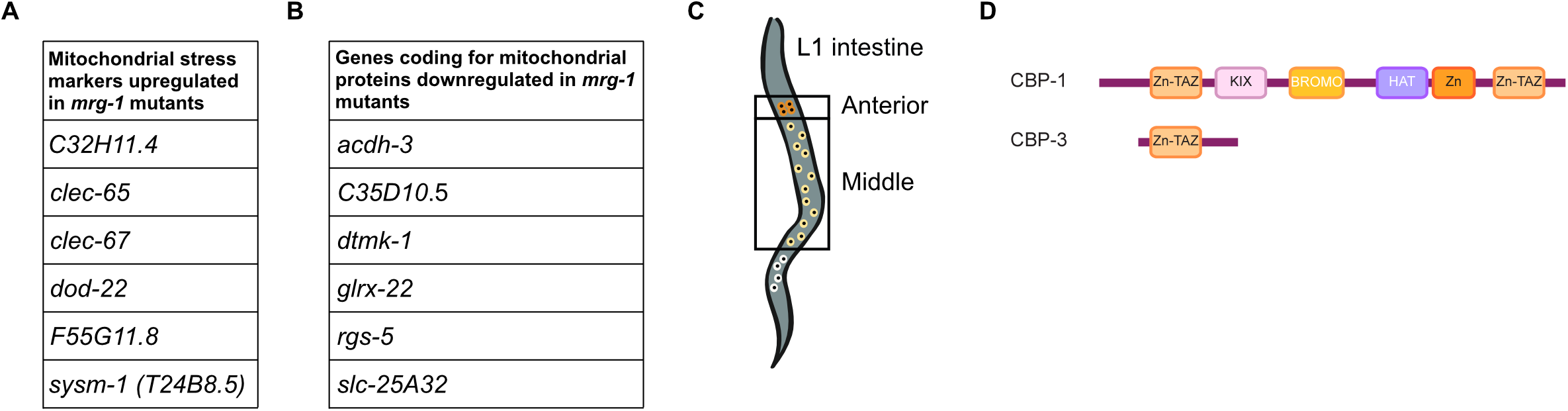
**A)** Mitochondrial stress markers from Campos et al., 2021 upregulated in *mrg-1* mutants. **B)** Genes coding for mitochondrial proteins downregulated in *mrg-1* mutants. **C)** Compartments in the *C. elegans* intestine. **D)** Schematic representation of CBP-1 and CBP-3 protein domains. A and B are from the analysis of the RNA-seq in Cabianca et al., 2019.

**Supplemental S2.**
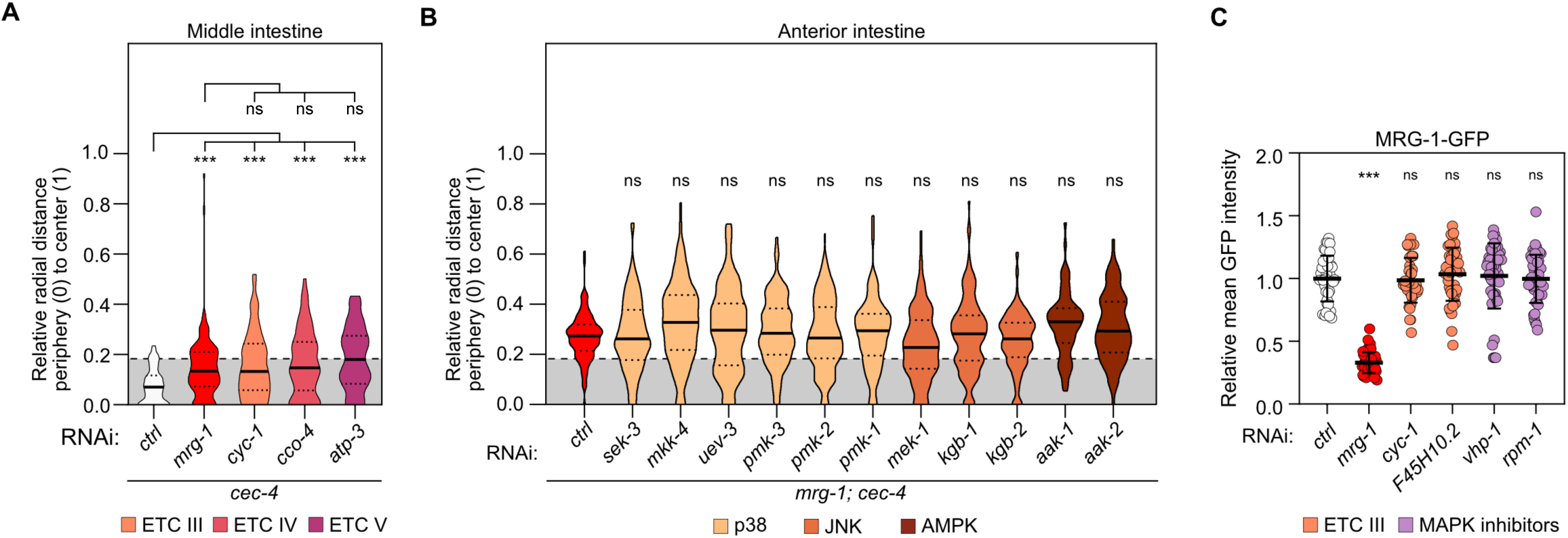
**A)** Relative radial distance distribution of the heterochromatin reporter foci in nuclei of anterior intestinal cells of *cec-4* mutant L1s exposed to the indicated RNAi. Results are from 3 biological replicates. **B)** Same as A), but in *mrg-1; cec-4* background. Results are from at least 2 biological replicates. **C)** Mean fluorescence intensity quantification of MRG-1-GFP, expressed from its endogenous locus (*ck45*(*mrg-1::degron::gfp*)) from images obtained by average intensity z-projections. Results are from 3 biological replicates, n (larvae) ≥ 42. For A, B and C, statistical significance was assessed using a Kruskal–Wallis non-parametric ANOVA. Pairwise comparisons were performed against the reference as indicated in each panel. *p< 0.05, ***p < 0.001, ns = not significant. For A and B, n (foci scored per condition) ≥ 125. Exact p-values and n (foci for A and B, worms for C, scored per condition) values are provided in Supplemental Table ST3.

**Supplemental S3.**
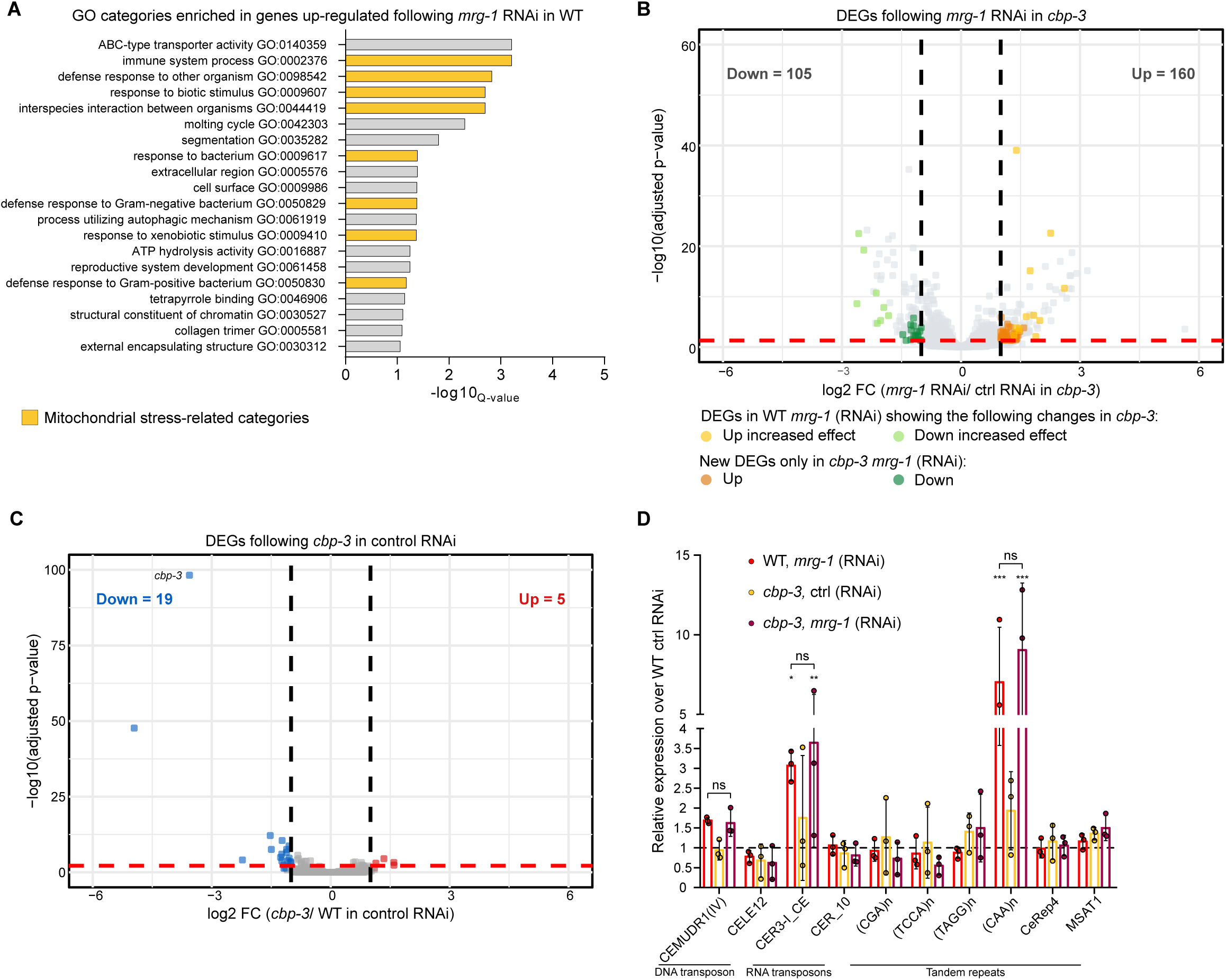
**A)** Gene Ontology terms enrichment of genes up-regulated by *mrg-1* RNAi in WT RNAseq. **B)** Volcano plot showing gene-expression changes following *mrg-1* RNAi relative to control RNAi in *cbp-3* mutants. Colored points indicate genes identified as differentially expressed following *mrg-1* RNAi in WT in (A) that were classified as increased effect in *cbp-3* mutants (see Methods for classification criteria), and genes differentially expressed only following *mrg-1* RNAi in *cbp-3*. **C)** Volcano plot showing genes differentially expressed in *cbp-3* mutants relative to WT, both in control RNAi. **D)** qPCR quantification of the relative expression of 10 repetitive elements in the indicated genotype and RNAi conditions, over WT animals treated with control RNAi. Results are from 3 biological replicates. Statistical significance was assessed using a two-way ANOVA, followed by Dunnett’s test for multiple comparisons to the WT control RNAi condition. *p< 0.05, ***p < 0.001, ns = not significant. Error bars represent the standard deviation. Exact p-values are provided in Supplemental Table ST3.

**Supplemental S4.**
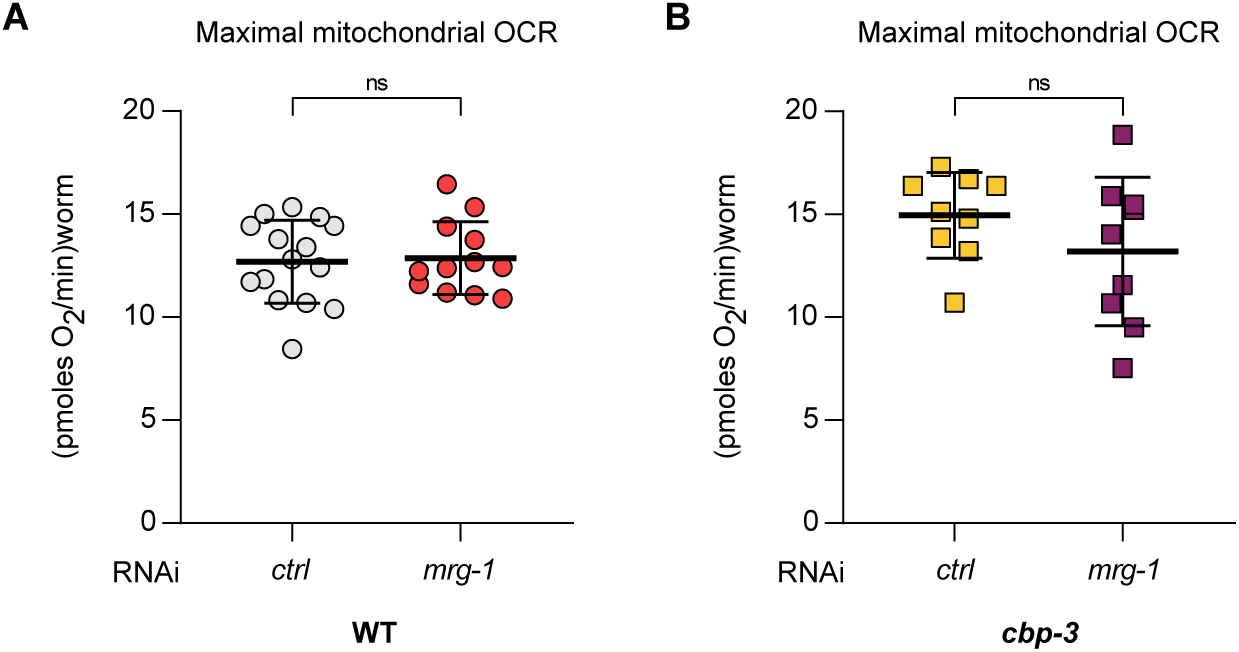
**A)** Maximal oxygen consumption rate (OCR) of WT adult worms exposed to *mrg-1* RNAi, measured by SeaHorse assay. Individual data points represent technical replicates from 5 independent biological replicates, 20-30 worms per technical replicate. **B)** Same as A), but in *cbp-3* mutants. Individual data points represent technical replicates from 3 independent biological replicates, 20-30 worms per technical replicate. For A and B, statistical significance was assessed using an unpaired t-test between the two conditions, ns = not significant. Exact p-values and n values are provided in Supplemental Table ST3.

